# A bipartite, low-affinity roadblock domain-containing GAP complex regulates bacterial front-rear polarity

**DOI:** 10.1101/2022.03.17.484758

**Authors:** Dobromir Szadkowski, Luís António Menezes Carreira, Lotte Søgaard-Andersen

## Abstract

The Ras-like GTPase MglA is a key regulator of front-rear polarity in rod-shaped *Myxococcus xanthus* cells. MglA-GTP localizes to the leading cell pole and stimulates assembly of the two motility machineries. MglA-GTP localization is spatially constrained by its cognate GEF, the RomR/RomX complex, and GAP, the MglB Roadblock-domain protein. RomR/RomX and MglB localize similarly with low and high concentrations at the leading and lagging poles, respectively. Yet, GEF activity dominates at the leading and GAP activity at the lagging pole by unknown mechanisms. Here, we identify RomY as a co-GAP that stimulates MglB GAP activity. The MglB/RomY interaction is low affinity, restricting complex formation to the lagging pole with the high MglB concentration. Our data support a model wherein RomY, by forming a low-affinity complex with MglB, ensures that MglB GAP activity is spatially precisely confined to the lagging pole, thereby constraining MglA-GTP to the leading pole establishing front-rear polarity.

## Introduction

Cell polarity enables essential cellular processes such as growth, division, differentiation, and motility ^1–3^. Small GTPases of the Ras superfamily are key cell polarity regulators in eukaryotes and bacteria ^4–8^, while they remain underexplored in archaea despite being abundant in several lineages ^9^. Typically, the function of small GTPases in cell polarity is coupled to their subcellular localization ^4–7^. A central unresolved question is how the precise subcellular localization of these GTPases is established.

Ras superfamily GTPases are molecular switches that alternate between an inactive, GDP-bound and an active, GTP-bound conformation ^10^. The nucleotide-dependent conformational changes center on the switch-1 and switch-2 regions close to the nucleotide-binding pocket, allowing the GTP-bound GTPase to interact with downstream effectors to implement a specific response ^10^. The activation/deactivation cycle is regulated by a cognate guanine-nucleotide exchange factor (GEF), which facilitates the exchange of GDP for GTP, and a GTPase activating protein (GAP), which stimulates the low intrinsic GTPase activity ^11,12^. Generally, the subcellular localization of a small GTPase is brought about by the localized activity of its cognate GEF, while the role played by its cognate GAP is less well-understood ^4,8^.

Motility in bacterium *Myxococcus xanthus* is an excellent model system to investigate how the spatiotemporal regulation of a small GTPase by its cognate GEF and GAP establishes dynamic cell polarity. *M. xanthus* cells are rod-shaped and translocate across surfaces with defined front-rear polarity, i.e. with a leading and lagging cell pole ^7,13^. In response to signaling by the Frz chemosensory system, front-rear polarity is inverted, and cells reverse their direction of movement ^14^. Motility and its regulation by the Frz system are prerequisites for multicellular morphogenesis with the formation of spreading, predatory colonies in the presence of nutrients and spore-filled fruiting bodies in the absence of nutrients ^7,13^. *M. xanthus* has two polarized motility systems. Gliding motility depends on the Agl/Glt complexes that assemble at the leading pole, adhere to the substratum, and disassemble at the lagging pole ^15,16^. In the type IV pili (T4P)-dependent motility system, T4P assemble at the leading pole ^17^ and undergo extension-adhesion-retraction cycles that pull a cell forward ^18,19^. Accordingly, during Frz-induced reversals, the cell pole at which the motility machineries assemble switches ^15,17,20^.

Front-rear polarity in *M. xanthus* is established by the so-called polarity module that consists of the small cytoplasmic GTPase MglA and its regulators. MglA generates the output of the polarity module and is essential for both motility systems ^21,22^. MglA follows the canonical scheme for small GTPases in cell polarity with the active GTP-bound state localizing to the leading cell pole, while the inactive MglA-GDP is diffuse in the cytoplasm ^23,24^. At the leading pole, MglA-GTP stimulates assembly of the Agl/Glt complexes ^16,25,26^ and extension of T4P ^27,28^ by interacting with downstream effectors. The cognate GEF and GAP of MglA control its nucleotide-bound state and localization. The RomR/RomX complex has MglA GEF activity ^29^. In this complex, RomX interacts with MglA to stimulate nucleotide exchange, and this activity is enhanced by RomR ^29^. Neither RomX nor RomR share homology with known GEFs in eukaryotes ^11,12,29^. MglB has MglA GAP activity *in vitro* ^23,24^. Structural analyses have demonstrated that MglB is a homodimeric Roadblock domain-containing protein and forms a 2:1 complex with MglA-GTP ^30–32^.

The RomR/RomX complex and MglB also localize polarly and, unexpectedly, localize in the same bipolar asymmetric pattern with a high concentration at the lagging and a low concentration at the leading pole ^23,24,29,33–35^. Nonetheless, *in vivo* evidence supports that GEF activity dominates at the leading pole ^29^, while GAP activity dominates at the lagging pole ^16,27,35,36^. RomR/RomX recruits MglA-GTP to the leading pole via two mechanisms: One depends on GEF activity, and in the second, the RomR/RomX complex interacts directly with MglA-GTP ^29^. MglB, via its GAP activity, excludes MglA from the lagging pole ^35,36^. Therefore, in the absence of MglB, MglA-GTP localizes more symmetrically at the cell poles ^23,24,36^, resulting in the formation of T4P at both poles ^27^ and lack of Agl/Glt complex disassembly at the lagging pole ^16^. Consequently, cells lose front-rear polarity, hyper-reverse erratically independently of the Frz system, and display little net movement. During the Frz-induced reversals, the polarity of MglA, MglB and RomR/RomX is inverted ^23,24,29,34^, thus, laying the foundation for the assembly of the motility machineries at the new leading pole. The mechanism underpinning the spatial separation of the GEF and GAP activities to the two cells poles is unclear.

Here, we investigated how the RomR/RomX GEF and MglB GAP activities are spatially separated. We report the identification of MXAN_5749 (from hereon RomY) and demonstrate that RomY is a co-GAP that stimulates MglB GAP activity. Notably, the MglB/RomY interaction is low affinity, and, therefore, MglB/RomY complex formation only occurs at the lagging cell pole with the high MglB concentration. Consequently, MglB GAP activity is stimulated only at the lagging pole, thereby restricting MglA-GTP to the leading pole. Thus, the key to precisely stimulating MglB GAP activity only at the lagging pole is that the MglB/RomY interaction is low-affinity.

## Results

### RomY is essential for the correct reversal frequency

Using a set of 1611 prokaryotic genomes, we previously used a phylogenomic approach to identify RomX ^29^. This approach was based on the observations that MglA and MglB homologs are widespread in prokaryotes ^37^. At the same time, RomR has a more narrow distribution and, generally, co-occurs with MglA and MglB ^33^. We, therefore, reasoned that proteins with a genomic distribution similar to RomR would be candidates for being components of the polarity module. Using this strategy, we also identified the uncharacterized protein RomY (encoded by MXAN_5749) (Fig. 1a).

**Fig. 1.**
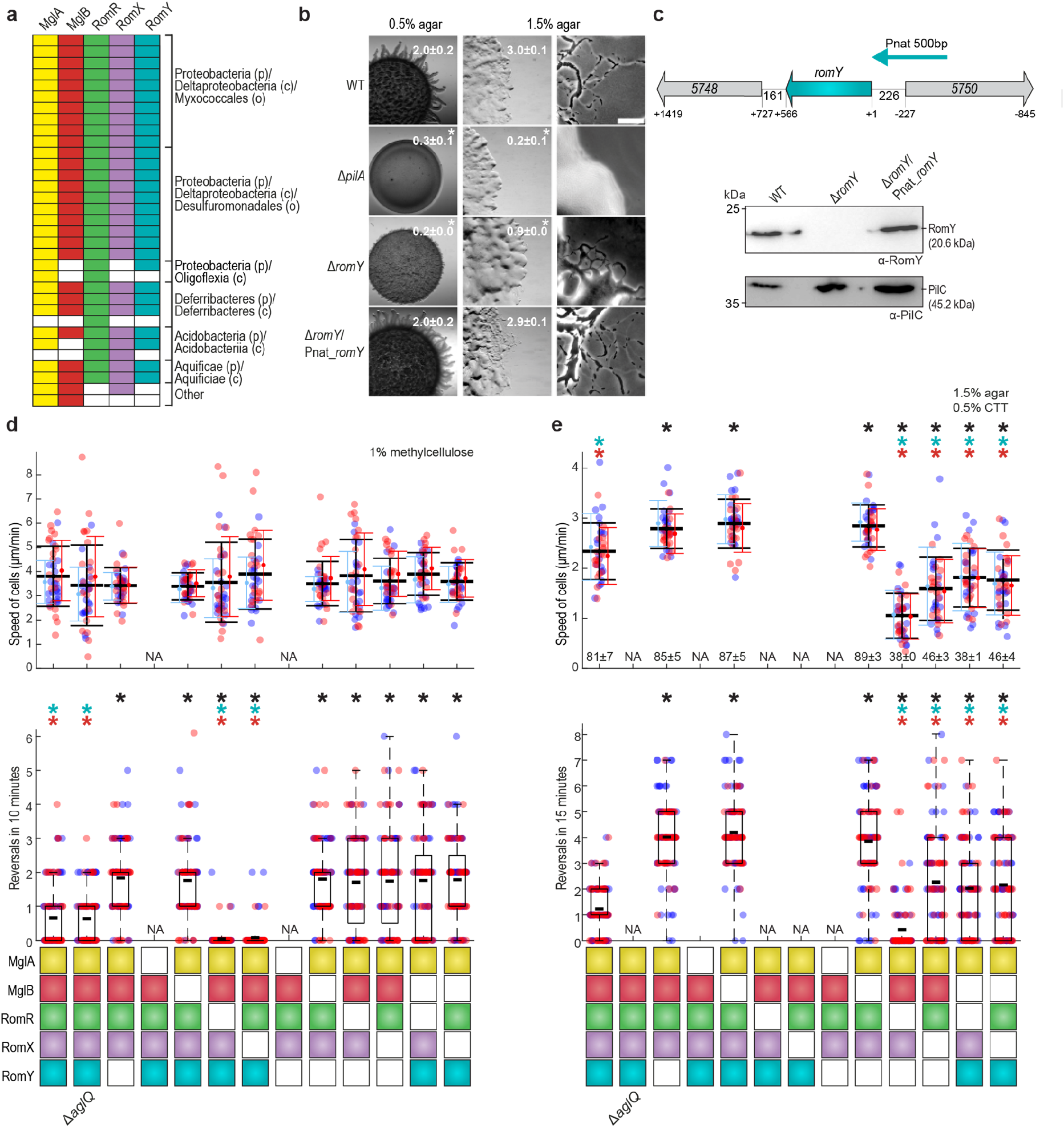
RomY is a component of the polarity module and important for correct reversals. **a.** RomY co-occurs with proteins of polarity module. Each column indicates the presence or absence of the relevant gene for the indicated proteins as colored or white boxes, respectively in a set of 1611 prokaryotic genomes. Lowest taxonomic level that includes all species in a group are indicated as phylum (p), class (c) and order (o). **b.** RomY is important for both motility systems. Cells were incubated on 0.5/1.5% agar with 0.5% CTT to score T4P-dependent/gliding motility. Scale bars, 1mm (left), 500 μm (middle), 50 μm (right). Numbers, colony expansion in mm in 24hrs as mean ± standard deviation (STDEV) (*n*=3); * *P*<0.05, two-sided Student’s *t*-test. **c.** *romY* locus and accumulation of RomY. Upper panel, *romY* locus; numbers in arrows, MXAN locus tags; numbers below, distance between stop and start codons. Cyan arrow, 500 bp fragment used for ectopic expression of *romY* and *romY-YFP*. Lower panel, immunoblot analysis of RomY accumulation. Cell lysates prepared from same number of cells were separated by SDS–PAGE and probed with *α*-RomY antibodies and *α*-PilC antibodies after stripping (loading control). The experiment was repeated twice with similar results. **d, e.** RomY is important for correct reversals. Boxes below diagrams indicate the presence or absence of indicated proteins as colored or white boxes, respectively. The Δ*aglQ* mutant is a control that T4P-dependent motility is scored in (**d**) and gliding in (**e**). Individual data points from two independent experiments with each *n*=20 cells (upper panels) and *n*=50 cells (lower panels) are plotted in red and blue. Upper diagrams, speed of cells moving by T4P-dependent motility (**d**) or gliding (**e**). Mean±STDEV is shown for each experiment and for both experiments (black). In (**e**), numbers indicate mean fraction±STDEV of moving cells. NA, not applicable because cells are non-motile. Lower panels, boxplots of reversals per cell in 10 or 15min; boxes enclose 25^th^ and 75^th^ percentiles, thick black line indicates the mean and whiskers the 10^th^ and 90^th^ percentiles. In all panels, * *P*<0.01, two-sided Student’s *t*-test. Black, cyan and red * indicate comparison to WT, the Δ*romY* strain and the Δ*mglB* strain, respectively. Source data for **b-e** are provided in Source Data file.

Based on sequence analysis, RomY is a 188-residue cytoplasmic protein. The RomY homologs identified in the 1611 genomes share a conserved N-terminal region, which includes residues 8-89 in RomY of *M. xanthus* and does not match characterized domain models, and a partially conserved C-terminal motif (Fig. S1a; Supplementary Table 1). The *romY* locus is partially conserved in Myxococcales, but none of the genes flanking *romY* has been implicated in motility (Fig. S1b).

To characterize RomY function, we generated a *romY* in-frame deletion mutation (Δ*romY*) in the wild-type (WT) strain DK1622. In population-based motility assays, cells were spotted on 0.5% and 1.5% agar that are favorable to T4P-dependent and gliding motility, respectively ^38^. On 0.5% agar, WT displayed long flares at the colony edge characteristic of T4P-dependent motility, while the Δ*pilA* mutant, which cannot assemble T4P, generated smooth colony edges; the Δ*romY* mutant formed shorter flares and had significantly reduced colony expansion compared to WT (Fig. 1b). On 1.5% agar, WT displayed single cells at the colony edge characteristic of gliding motility, while the Δ*aglQ* mutant, which lacks a component of the Agl/Glt machinery, did not. The Δ*romY* mutant had fewer single cells at the colony edge and significantly reduced colony expansion compared to WT (Fig. 1b). In complementation experiments, ectopic expression of *romY* from its native promoter on a plasmid integrated in a single copy at the Mx8 *attB* site restored the defects in both motility systems (Fig. 1b,c). Ectopically produced RomY accumulated at a level similar to that in WT (Fig. 1c).

Using assays to monitor the motility characteristics with single-cell resolution, we observed that for both motility systems, Δ*romY* cells moved with speeds similar to WT (Fig. 1d,e, upper panels), but reversed at a significantly higher frequency than WT (Fig. 1d,e, lower panels).

Finally, in the absence of the FrzE kinase, which is essential for Frz-induced reversals ^39^, Δ*romY* cells still hyper-reversed (Fig. S2). We conclude that RomY is not necessary for motility *per se* but for maintaining the correct reversal frequency. Moreover, the epistasis experiment support that RomY acts downstream of the Frz system to maintain the correct reversal frequency.

### A Δ*romY* mutant has the same phenotype as the Δ*mglB* mutant

We performed epistasis tests using single-cell motility characteristics as readouts to test whether RomY functions in the same genetic pathway as MglA, MglB, RomR and RomX. In T4P-dependent motility (Fig. 1d), all single and double mutants except for the Δ*mglA* and the Δ*mglA*Δ*romY* mutants, none of which displayed movement, had speeds similar to WT. The Δ*mglB*, Δ*romY* and Δ*mglB*Δ*romY* mutants had the same hyper-reversing phenotype. As previously reported, the Δ*romR* and Δ*romX* mutants hypo-reversed ^29,40^, while the Δ*romR*Δ*romY* and Δ*romX*Δ*romY* double mutants had reversal phenotypes similar to that of the Δ*romY* mutant. Because the Δ*mglB*, Δ*romY* and Δ*mglB*Δ*romY* mutants have the same hyper-reversal phenotype, we included the Δ*mglB*Δ*romR* and Δ*mglB*Δ*romX* double mutants; these two mutants had reversal phenotypes similar to that of the Δ*mglB* mutant and the Δ*romR*Δ*romY* and Δ*romX*Δ*romY* double mutants.

In gliding motility (Fig. 1e), the Δ*mglA*, Δ*romR* and Δ*romX* mutants are non-motile because no or insufficient MglA-GTP accumulate to stimulate Agl/Glt complex formation. Again, the Δ*mglB*, Δ*romY* and Δ*mglB*Δ*romY* mutants were similar with respect to speed and had the same hyper-reversal phenotype. Notably, the Δ*romY* mutation, similarly to the Δ*mglB* mutation ^29^, partially alleviated the deleterious effect of the Δ*romR* and Δ*romX* mutations on gliding.

The epistasis experiments support that RomY acts in the same pathway as MglA, MglB, RomR and RomX. Moreover, we conclude that RomY, similarly to MglB (Fig. S2) ^16,23,24,27^, acts downstream of the Frz system to maintain correct reversals. Notably, lack of MglB or RomY causes strikingly similar phenotypes with (1) Frz-independent hyper-reversals and (2) partial suppression of the gliding motility defect in the Δ*romR* and Δ*romX* mutants. The Frz-independent hyper-reversals caused by lack of MglB result from the accumulation of MglA-GTP at both poles with the concomitant loss of front-rear polarity ^16,33,35^ The defect in gliding motility in the Δ*romR* and Δ*romX* mutants is caused by a lack of MglA GEF activity and, therefore, a low MglA-GTP level. These considerations support that lack of RomY, similarly to lack of MglB, causes increased accumulation of MglA-GTP.

At least three non-mutually exclusive scenarios can explain the increased accumulation of MglA-GTP in the Δ*romY* mutant: RomY (1) inhibits RomR/RomX GEF activity, (2) stimulates MglB GAP activity, or (3) has MglA GAP activity. These scenarios make different predictions. In scenario (1), the Δ*romY* mutation would not suppress the gliding defect in the Δ*romR* and Δ*romX* mutants, and the effects of the Δ*mglB* and Δ*romY* mutations on reversal frequency would be additive. Scenario (2) and (3) predict that the Δ*romY* mutation would suppress the gliding defect in the Δ*romR* and Δ*romX* mutants; however, in scenario (2), the effects of the Δ*mglB* and Δ*romY* mutations would not be additive, while they would be additive in scenario (3). All the results of the epistasis experiments agree with scenario (2), supporting that RomY stimulates MglB GAP activity.

### RomY is an MglB co-GAP *in vitro*

Prompted by the above considerations, we examined the effect of RomY on MglA GTPase activity *in vitro* by measuring released GDP in a coupled enzyme assay or released phosphate (P_i_) in a malachite green-based assay using purified MglA-His_6_, His_6_-MglB and Strep-RomY (Fig. S3a). MglA-His_6_ was preloaded with GTP and then mixed with equimolar amounts of MglB-His_6_ and/or Strep-RomY in the presence of ~300-fold molar excess of GTP. MglA-His_6_ alone had a low GTPase activity (Fig. 2a). His_6_-MglB stimulated MglA-His_6_ GTPase activity, while Strep-RomY did not. Importantly, in the presence of both MglB-His_6_ and Strep-RomY, MglA-His_6_ GTPase activity increased two-fold compared to MglB-His_6_ only in both assays. Neither MglB-His_6_ nor Strep-RomY had GTPase activity. In conclusion, RomY stimulates MglA GTPase activity but only in the presence of MglB.

**Fig. 2.**
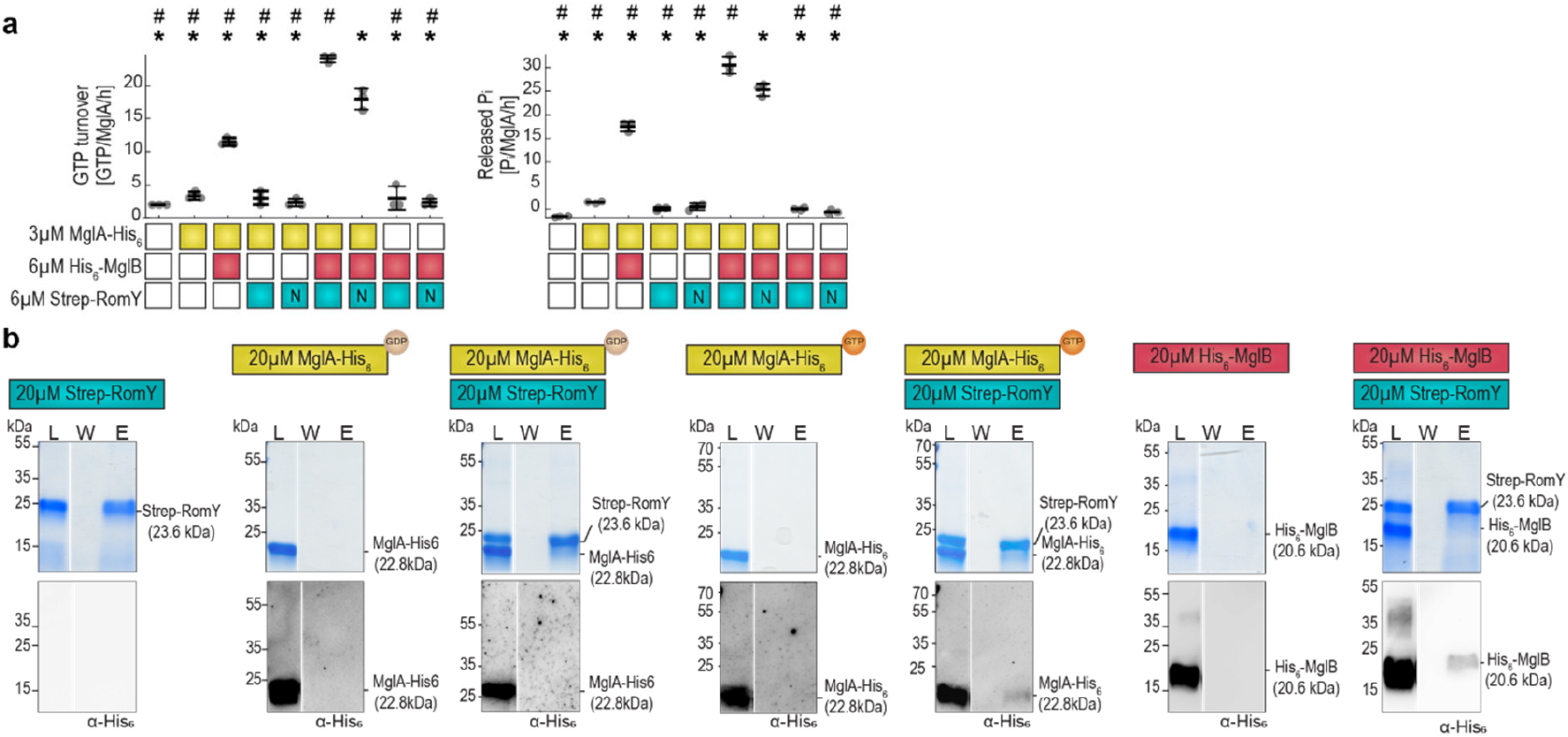
RomY stimulates MglA GTPase activity in the presence of MglB and interact with MglA-GTP and MglB. **a.** RomY stimulates MglA GTPase activity in the presence of MglB. GTPase activity measured as GTP turnover in enzymatic coupled regenerative GTPase assay (left) and released inorganic phosphate in malachite green assay (right), after 1h of incubation. Boxes below diagrams indicate the presence or absence of indicated proteins as colored or white boxes, respectively, GTP was added to 1mM. For Strep-RomY, N indicate Strep-RomY^N^. Individual data points from three independent experiments are in gray and mean±STDEV indicated. * and #, *P*<0.05, two-sided Student’s *t*-test with samples compared to MglA-His_6_/His_6_-MglB/Strep-RomY and MglA-His_6_/His_6_-MglB/Strep-RomY^N^, respectively. **b.** RomY interacts with MglB and MglA-GTP. Proteins were mixed with final concentrations and 10mM GTP/GDP as indicated in the schematics for 30min at RT, DSP added (final concentration 200μM, 5min, RT), DSP quenched, and proteins applied to Strep-Tactin coated magnetic beads. Fractions before loading (L), the last wash (W) and after elution (E) were separated by SDS–PAGE, gels stained with Coomassie Brilliant Blue (upper panels) and subsequently probed with α-His_6_ antibodies (lower panels). All samples were treated with loading buffer containing 100mM DTT to break crosslinks before SDS-PAGE. For each combination, fractions were separated on the same gel. Gaps between lanes indicate lanes deleted for presentation purposes. The experiments in **b** were repeated twice with similar results. Source data for **a**-**b** are provided in Source Data file.

Next, we investigated how MglA, MglB and RomY interact using pull-down experiments with Strep-RomY as the bait. Strep-RomY alone bound to Strep-Tactin beads, while His_6_-MglB, MglA-His_6_-GTP, MglA-His_6_-GDP and the His_6_-MalE negative control neither bound alone nor in the presence of Strep-RomY (Fig. S3a,b). We, therefore, speculated that the interaction(s) between RomY and MglB and/or MglA could be low affinity resulting in transient complex formation. To test this possibility, we added the protein cross-linked dithiobis(succinimidyl propionate) (DSP) to the protein mixtures before affinity chromatography. After elution, crosslinks were broken with dithiothreitol (DTT) and proteins separated by SDS-PAGE. Cross-linked Strep-RomY bound to the Strep-Tactin beads (Fig. 2b), while neither cross-linked His_6_-MalE, which served as a negative control, MglB-His_6_ nor MglA-His_6_ preloaded with GTP or GDP did (Fig. 2b; Fig. S3c). However, after crosslinking in the presence of Strep-RomY protein, His_6_-MglB and MglA-His_6_ preloaded with GTP were retained, while MglA preloaded with GDP and His_6_-MalE were not (Fig. 2b; Fig. S3c).

We conclude that RomY interacts separately with MglB and MglA-GTP. Because RomY stimulates MglA GTPase activity in an MglB-dependent manner, we conclude that the three proteins also form a complex in which all three proteins are present. Because RomY alone does not have MglA GAP activity even though the two proteins interact, we refer to RomY as an MglB co-GAP. The observation that all interactions were only observed after crosslinking suggests that they are low affinity giving rise to transient complex formation.

To gain insights into how RomY may interact with MglA-GTP and the MglB homodimer, we generated a structural model of RomY using AlphaFold ^41^ and the ColabFold pipeline ^42^. In all five models generated, residues 7-90, which covers the N-terminal conserved region from residue 8-89 (Fig. S1a), were predicted to fold into a globular domain with high confidence based on Predicted Local Distance Difference Test (pLDDT) and predicted alignment error (pAE), while the remaining parts of RomY was modelled with lower confidence (Fig. 3a; Fig. S4a). Because the N-terminal conserved part of RomY extends from residue 8 to 89, from hereon we refer to residue 1-89 as the N-terminal domain of RomY.

**Fig. 3.**
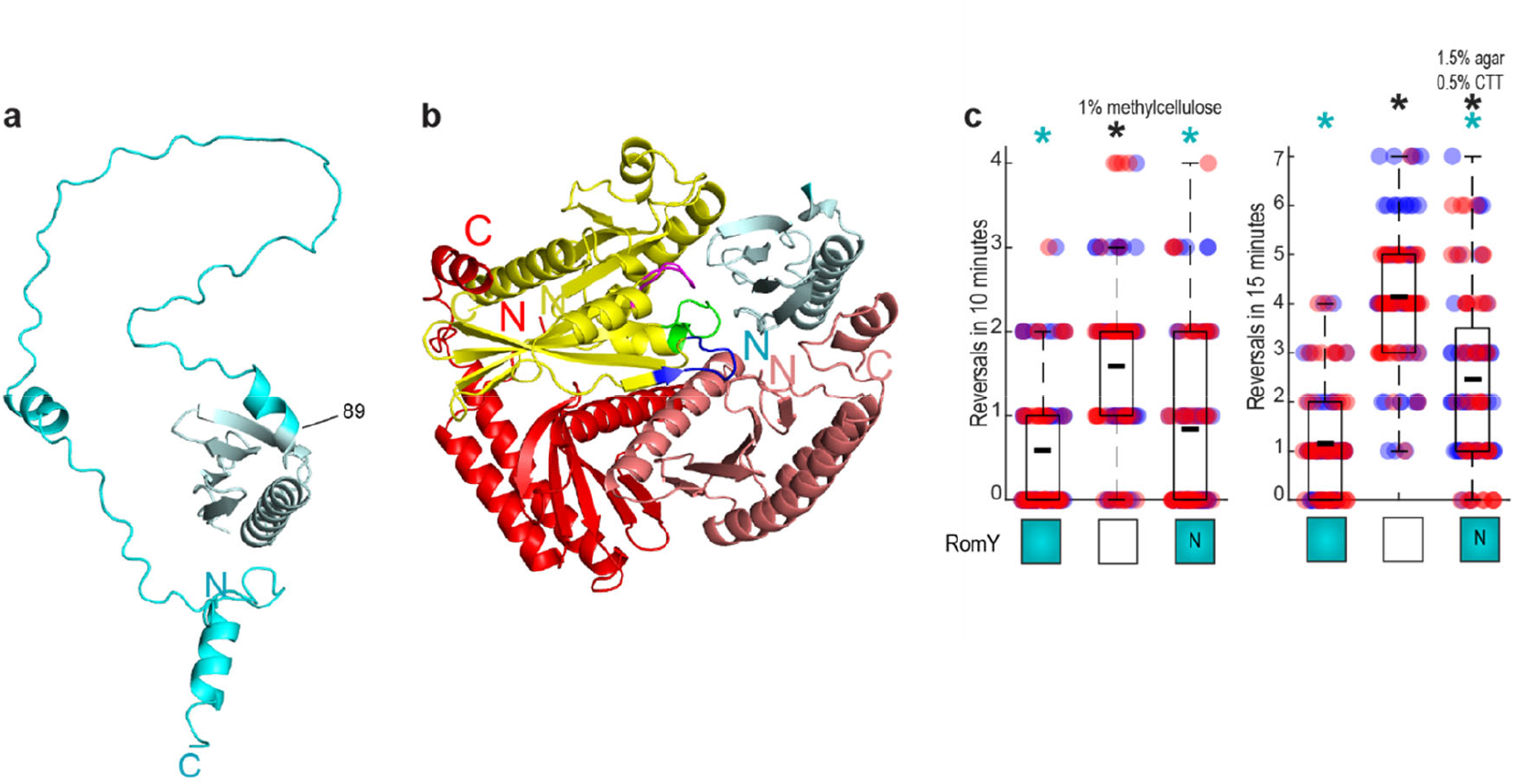
The N-terminal domain of RomY has partial RomY activity. **a.** AlphaFold model of RomY. RomY was modeled as a monomer. The N-terminal conserved region up to residue 89 is in teal and the remaining part in cyan. Model rank 1 is shown. **b.** AlphaFold-Multimer model of the MglA:(MglB)_2_:RomY complex. The MglA monomer is in yellow and with the P-loop in purple, switch region-1 in blue and switch region-2 in green, the MglB homodimer in red, and the N-terminal domain of RomY in teal. **c**. The N-terminal domain of RomY has partial RomY activity. Reversals were tracked in single cells for T4P-dependent and gliding motility as in Fig. 1**d**, **e**. Boxes below diagrams indicate the presence or absence of RomY as colored or white boxes, respectively. N indicates RomY^N^. Individual data points from two independent experiment with each *n*=50 cells are plotted in red and blue. Boxplot is as in Fig. 1**d**, **e**. * *P*<0.05, two-sided Student’s *t-* test with comparison to WT (black) and the Δ*romY* mutant (cyan). Source data for **c** is provided in Source Data file.

To understand how RomY may interact with MglA, the MglB homodimer and the two proteins in parallel, we used AlphaFold-Multimer ^43^ to generate models of MglA:RomY, (MglB)_2_:RomY and MglA:(MglB)_2_:RomY complexes as well as of an MglA:(MglB)_2_ complex. All five models of MglA:(MglB)_2_ were predicted with high confidence and are in overall agreement with the solved structure of MglA-GTPγS:(MglB)_2_ ^30,32^ (Fig. S4a,b) documenting the quality of the prediction and that AlphaFold-Multimer models MglA in the GTP-bound form in this complex.

For each of the five models of MglA:RomY, (MglB)_2_:RomY and MglA:(MglB)_2_:RomY complexes, we obtained high confidence predictions based on pLDDT and pAE scores including residues 7-90 in RomY (Fig. S4a). Therefore, we only considered the N-terminal domain of RomY in the models. In the MglA:RomY model, MglA had a structure similar to that of the solved structure of MglA-GTPγS ^30,32^ (Fig. S4c), and, thus, AlphaFold-Multimer models MglA in the GTP bound form. Notably, the N-terminal domain of RomY associated with MglA close to the nucleotide-binding pocket (Fig. S4c). In the (MglB)_2_:RomY model, the MglB homodimer was similar to the solved structure ^30,32^, and the RomY N-terminal domain interacted asymmetrically with the two MglB monomers using a different surface than for the interaction with MglA (Fig. S4d). Finally, in the model of all three proteins, the MglA:(MglB)_2_ part was similar to the solved structure of the MglA-GTPγS:(MglB)_2_ complex ^30,32^ (Fig. S4e), documenting that AlphaFold-Multimer models MglA in the GTP bond state. Importantly, the N-terminal domain of RomY interacted with MglA and one of the MglB monomers in the MglB homodimer, and was positioned close to the nucleotide-binding pocket of MglA (Fig. 3b). Thus, the MglA:(MglB)_2_:RomY model supports that all three proteins interact to form a complex in which they interact in all three pairwise directions and in which MglA is in the GTP-bound state. Also, this model agrees with the interactions detected in the pull-down experiments (Fig. 2b). Moreover, this model suggests that the N-terminal domain of RomY has a key role in the co-GAP activity of RomY.

To test this structural model, we generated a RomY variant (RomY^N^) that was truncated for the C-terminal part of RomY and only included residue 1-89 (Fig. 3a-b; Fig. S1a). In the GTPase assays, RomY^N^ stimulated MglA GTPase activity in the presence of MglB almost as efficiently as full-length RomY (Fig. 2a; Fig. S3a). *In vivo*, a mutant synthesizing RomY^N^ as the only RomY protein had a motility phenotype between WT and the Δ*romY* mutant (Fig. 3c; Fig. S5a,b). Although neither purified Strep-RomY^N^ nor RomY^N^ synthesized *in vivo* was detectable in immunoblot analysis with α-RomY antibodies, these observations support that the N-terminal domain of RomY interacts with MglA-GTP as well as the MglB dimer and that the co-GAP activity largely resides in this region of RomY.

### RomY is essential for sufficient MglB GAP activity *in vivo*

MglB alone has MglA GAP activity *in vitro*. However, the Δ*mglB* and Δ*romY* mutations cause similar motility defects *in vivo* suggesting that RomY is required for sufficient MglB GAP activity *in vivo*. Alternatively, MglB alone is sufficient for GAP activity *in vivo*, but its concentration is too low to stimulate MglA GTPase activity efficiently. To resolve the importance of RomY for MglB GAP activity *in vivo*, we overexpressed MglB in the presence or absence of RomY. Cells with a low level of MglA-GTP are non-motile by gliding but move with WT speed and a reduced reversal frequency using the T4P-dependent motility system ^27,29,40^ (Cf. Fig. 1d,e, Δ*romR* and Δ*romX* mutants). Therefore, we used speed and reversal frequency in T4P-dependent motility as precise and sensitive readouts of GAP activity in these experiments.

MglB and RomY accumulated independently when expressed from their native loci (Fig. 4a). MglB was ectopically overproduced using a vanillate-inducible promoter (P_van_) in Δ*mglB* strains. In the absence of vanillate, MglB was not detectable in immunoblots; upon addition of 500 μM vanillate, MglB accumulated at an ~20-fold higher level than in WT and independently of RomY. The level of RomY was unaffected by the increased MglB accumulation (Fig. 4a). In the absence of vanillate, the Δ*mglB*/P_van__*mglB* strain containing RomY was similar to the Δ*mglB* strain and hyper-reversed (Fig. 4b; Fig. S6). Importantly, in the presence of vanillate, cells of this strain moved with WT speed but had a reversal frequency significantly below that of WT (Fig. 4b; Fig. S6), indicating a low MglA-GTP concentration. By contrast, in the absence of RomY, MglB overproduction did not affect the reversal frequency. In the inverse experiments, RomY was ectopically overproduced from a vanillate-inducible promoter in the absence or presence of MglB (Fig. 4a; Fig. S6). RomY overproduction (~10-fold higher than the RomY level in WT) in the presence of MglB resulted in a reversal frequency significantly below that of WT. By contrast, RomY overexpression in the absence of MglB did not affect the reversal frequency.

**Fig. 4.**
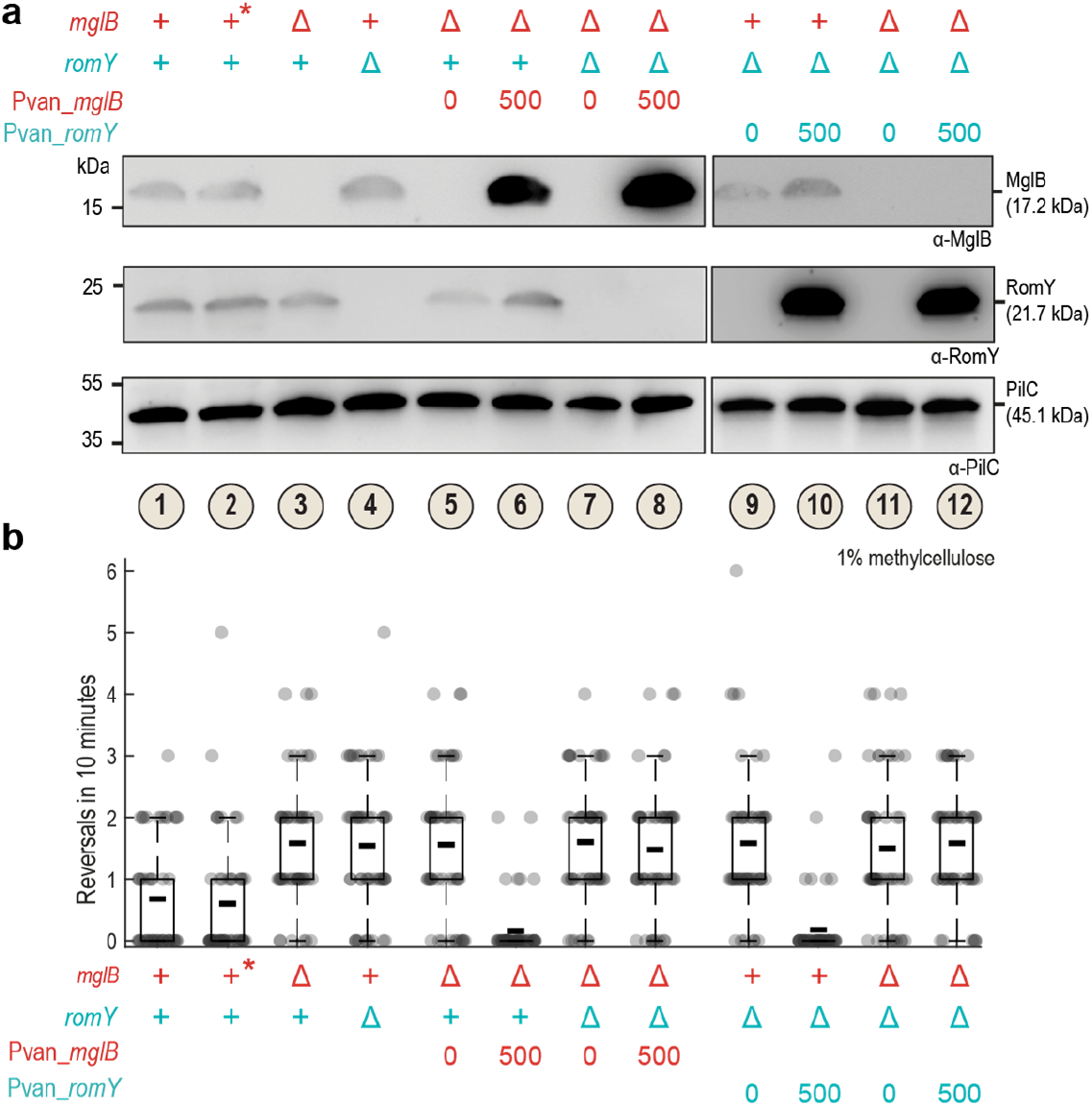
RomY is essential for sufficient MglB sufficient GAP activity *in vivo*. **a.** Analysis of MglB and RomY accumulation by immunoblot analysis in induction experiments. Strains of the indicated genotypes were grown in the presence and absence of 500μM vanillate for 3h as indicated. Cell lysates prepared from the same number of cells for each sample were separated by SDS–PAGE and probed sequentially with α-MglB, α-RomY and α-PilC (loading control) antibodies with stripping of the membrane before the second and third antibodies. In the legend, + indicates presence of WT gene, Δ in-frame deletion, 0/500μM vanillate concentration, and * the WT grown in the presence of 500μM vanillate. Samples 1-8 and 9-12 were separated on different SDS-PAGE gels that both contained samples 1-4 to enable comparisons of samples between different gels. The experiment was repeated twice with overall similar results. **b**. Analysis of reversals in T4P-dependent motility upon overproduction of MglB or RomY. Cells were treated as in (**a**) and then T4P-dependent single cell motility analyzed. Legend is as in (**a**). Individual data points from a representative experiment with *n*=50 cells are plotted in gray. Because the experiment relies on induction of gene expression, protein levels vary slightly between experiments, making the direct comparison between biological replicates difficult. Consequently, data from only one representative experiment is shown. Boxplots are as in Fig. 1**d**. The experiment was repeated twice with overall similar results. Source data for **a-b** are provided in Source Data file.

We conclude that MglB and RomY accumulate independently of each other. In addition, neither MglB nor RomY alone, even when highly overproduced, is sufficient to stimulate sufficiently MglA GTPase activity *in vivo*. Rather MglB, even when overproduced, depends on RomY for GAP activity *in vivo;* similarly, RomY only results in GAP activity in the presence of MglB *in vivo*. These observations also support that RomY is an MglB co-GAP and essential for sufficient MglB GAP activity *in vivo*. Notably, overproduction of either MglB or RomY in the presence of WT levels of RomY or MglB, respectively, results in increased GAP activity compared to WT. These observations corroborate that the rate-limiting step for MglB/RomY GAP activity is complex formation and not protein concentration, thus, supporting that the MglB/RomY interaction is low affinity.

### RomY localizes dynamically to the lagging cell pole in an MglB-dependent manner

Our data are consistent with RomY acting as a co-GAP for MglB *in vitro* and *in vivo*. To resolve if RomY contributes to the spatial regulation of MglB GAP activity *in vivo*, we determined RomY localization using an ectopically expressed, active RomY-YFP fusion (Fig. S7a,b). By snapshot analysis, RomY-YFP localized in a highly asymmetric pattern with 81% of cells having unipolar or asymmetric bipolar localization (Fig. 5a). In moving cells, the large cluster localized highly asymmetrically to the lagging cell pole (Fig. 5b; see also below). Moreover, RomY-YFP localization was dynamic, and after a reversal, RomY-YFP localized to the new lagging pole (Fig. 5b).

**Fig. 5.**
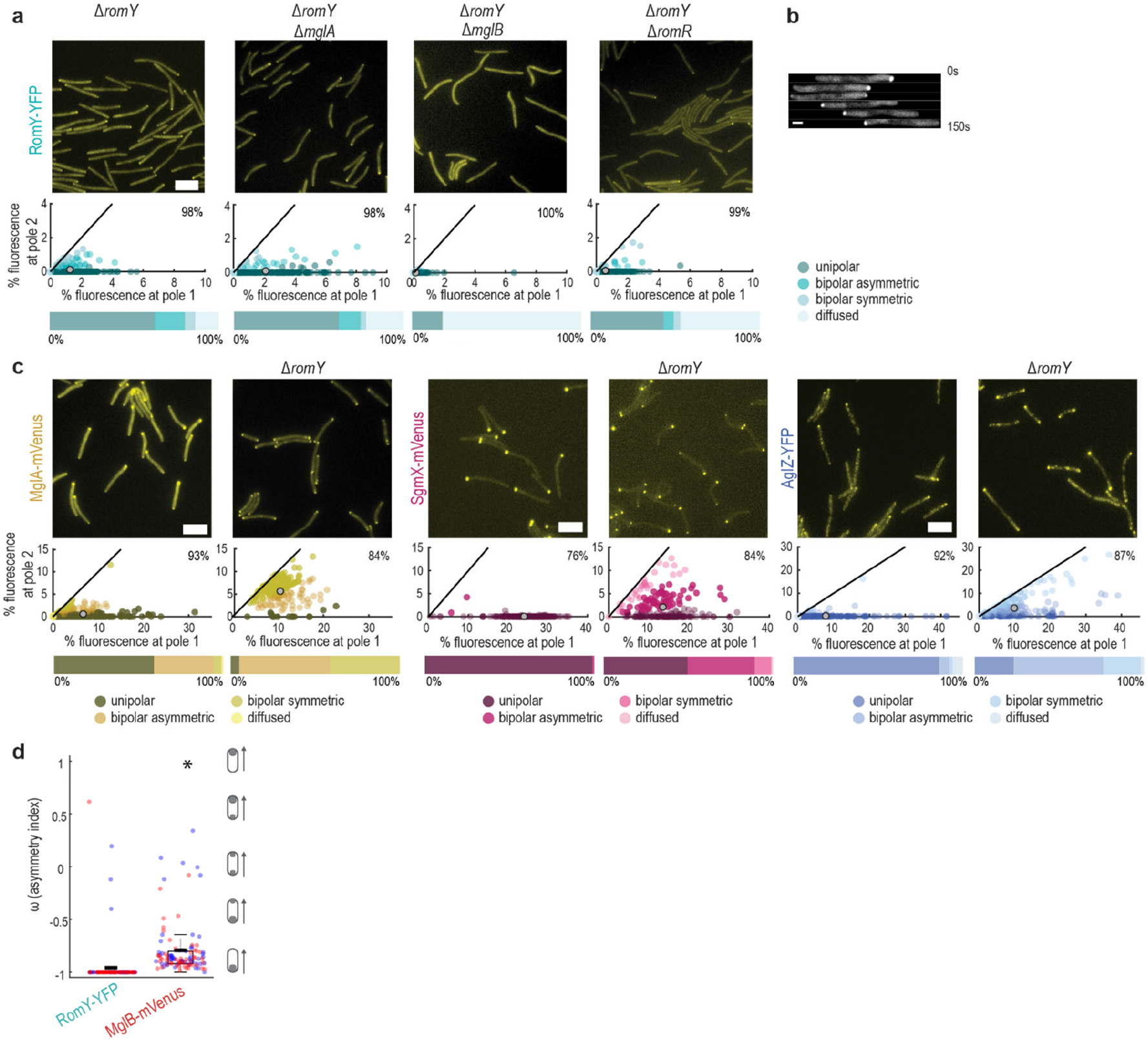
RomY localizes dynamically to the lagging cell pole. **a.** RomY-YFP localization by epi-fluorescence microscopy. In the scatter plot, the percentage of total fluorescence at pole 2 is plotted against the percentage of total fluorescence at pole 1 for all cells with polar cluster(s). Pole 1 is per definition the pole with the highest fluorescence. Individual cells are color-coded according to its localization pattern. Black lines are symmetry lines, grey spots show the mean and numbers in the upper right corner the mean percentage of total fluorescence in the cytoplasm. Horizontal bars below show the percentage of cells with a polar localization pattern and diffuse localization according to the color code. *n*=200 cells in all strains. Scale bar, 5 μm. **b.** RomY-YFP is dynamically localized to the lagging pole. Cells were imaged by time-lapse epi-fluorescence microscopy every 30s. Scale bar, 1*μ*m. **c.** MglA-mVenus, Sgmx-mVenus and AglZ-YFP localization in the absence of RomY. Cells were imaged by epi-fluorescence microscopy, scatter plots and percentage of cells with a particular localization pattern were determined as in **a**. *n* = 200 cells for all strains. Scale bar, 5 μm. **d.** Comparison of RomY-YFP and MglB-mVenus asymmetry in moving cells. Cells were imaged by time-lapse epi-fluorescence microscopy every 30s. An asymmetry index (ω) was calculated for cells that moved for three or more successive frames without reversing and excluding the first frame after a reversal and the last frame before a reversal (see Methods). As indicated in the schematics, ω = −1, unipolar at lagging pole, ω = +1, unipolar at leading pole, and ω = 0, bipolar symmetric. Individual data points from two independent experiments (27/33 cells and 48/54 data points for RomY-YFP, and 7/11 cells and 53/53 data points for MglB-mVenus) are plotted in red and blue. Boxplot is as in Fig. 1**d, e**. * *P*<0.005, two-sided Student’s *t*-test. Experiments in **a-c** were repeated twice with similar results. Source data for **a, c-d** are provided in Source Data file.

To determine how RomY-YFP is targeted to the lagging pole, we interrogated RomY-YFP localization in the absence of MglA, MglB or RomR, which we used as a proxy for the RomR/RomX complex (Fig 5a). RomY-YFP accumulated as in WT in the absence of each of these proteins (Fig. S7b). In the absence of MglA, the total polar RomY-YFP signal was as in WT and the protein was slightly more unipolar (Fig. 5a). In the absence of MglB, RomY-YFP polar localization was strongly reduced, and 81% of cells had no polar signals. In the absence of RomR, RomY-YFP polar localization was also decreased, but 54% of cells still had a weak polar signal. Polar localization of MglB, MglA and RomX is strongly reduced or even abolished in the absence of RomR ^29,33,35,36^, but MglA, RomR and RomX are still polarly localized in the absence of MglB ^36^. In agreement with the direct interaction between RomY and MglB, we, therefore, conclude that MglB is the primary polar targeting determinant of RomY-YFP, while MglA has at most a minor role in polar RomY localization. We note that in the absence of MglB, there is still some residual unipolar RomY-YFP localization. MglA is more bipolarly localized in the absence of MglB, supporting that MglA does not bring about this residual polar localization and that yet to be identified factor(s) may have a role in RomY polar localization.

To determine the effect of RomY on the proteins of the polarity module, we focused on MglA because this protein generates the output of this module. We also analyzed the two MglA-GTP effectors SgmX and AglZ that localize to the leading pole in an MglA-dependent manner to stimulate the formation of T4P ^27,28^ and assembly of the Agl/Glt complexes ^15^, respectively and, thus, provide a functional readout of the state of the polarity module. As previously shown, MglA-mVenus localized in a highly asymmetric pattern in WT (Fig. 5c; Fig. S7c). Notably, in the absence of RomY, polar localization of MglA-mVenus was increased and switched toward more bipolar symmetric (Fig. 5c). Thus, RomY, as previously observed for MglB, is required to exclude MglA-GTP from the lagging pole. In agreement with these observations and that the Δ*romY* mutant hyper-reverses in a Frz-independent manner, SgmX-mVenus and AglZ-YFP were shifted from strongly unipolar toward bipolar symmetric in the absence of RomY (Fig. 5c; Fig. S7d,e). Importantly, in the Δ*mglB* mutant, localization of SgmX and AglZ is also shifted toward bipolar symmetric ^27,35^.

Altogether, we conclude that MglB is the primary determinant of polar RomY localization, that RomY stimulates MglB GAP activity at the lagging pole, and, together with MglB, RomY is required to exclude MglA-GTP from this pole.

### RomY specifically stimulates MglB GAP activity at the lagging cell pole

Because MglB is bipolarly asymmetrically localized, we reasoned that for RomY to only stimulate MglB activity at the lagging pole and, thus, spatially confine RomY/MglB GAP activity to this pole, RomY would have to be more asymmetrically localized to the lagging pole than MglB. To this end, we determined the polar asymmetry of RomY-YFP and an active MglB-mVenus fusion (Fig. S7a,f) in moving cells. As shown in Fig. 5d, RomY-YFP was almost exclusively unipolar, while MglB-mVenus was almost exclusively bipolar asymmetric. Thus, RomY is significantly more asymmetrically localized to the lagging pole than MglB supporting that RomY only stimulates MglB GAP activity at this pole.

## Discussion

In the rod-shaped *M. xanthus* cells, the activity of the small GTPase MglA is spatially restricted to the leading cell pole by the joint action of its cognate GEF and GAP. This spatial regulation ensures that the two motility systems only assemble at this cell pole and is, thus, critical for directed motility. The RomR/RomX GEF and the MglB GAP localize similarly to the two poles but with GEF and GAP activity dominating at the leading and lagging cell pole, respectively. How this spatial separation of these two activities is brought about has remained unknown. Here, we report the identification of the previously uncharacterized RomY protein and demonstrate that it is an integral part of the polarity module. Specifically, RomY functions as an MglB co-GAP and, by forming a low-affinity complex with MglB, precisely stimulates and restricts MglB GAP activity to the lagging cell pole.

*In vitro* MglB alone has MglA GAP activity, while RomY alone does not. However, RomY stimulates MglA GTPase activity *in vitro* in the presence of MglB. *In vitro* RomY interacts independently with MglB and MglA-GTP in pull-down experiments. Because RomY interacts separately with MglB and MglA-GTP but does not have GAP activity on its own, we refer to RomY as an MglB co-GAP. Canonical GAPs of Ras-like GTPases supply either an arginine finger or an asparagine thumb to complete the active site of the GTPase ^11,12^. By contrast, the MglB dimer interacts asymmetrically with MglA-GTP and brings about the repositioning of amino acid residues in MglA to generate the active site for GTP hydrolysis ^30–32^. An AlphaFold-Multimer based structural model of the MglA:(MglB)_2_:RomY complex supports that the N-terminal domain of RomY interacts with one of the MglB monomers in the MglB homodimer as well as with MglA close to the nucleotide-binding pocket. Consistently, this RomY domain has partial co-GAP activity *in vitro* and partial RomY activity *in vivo*. Continued biochemical work and structural studies will be required to decipher the exact mechanism by which RomY functions as a co-GAP. Nonetheless, we speculate that RomY could increase the affinity of the MglB homodimer for MglA-GTP and/or cause conformational changes in the active site of MglA.

*In vitro* MglB alone has GAP activity, which under the conditions of the enzyme assay is stimulated ~two-fold by RomY. However, the Δ*romY* mutant phenocopies the Δ*mglB* mutant supporting that RomY is essential for sufficient MglB GAP activity to regulate polarity *in vivo*. Even when MglB was overproduced 20-fold, its activity was dependent on RomY. Our data do not allow us to distinguish whether RomY is essential for MglB GAP activity or for *sufficiently high* MglB GAP activity *in vivo* for MglB to regulate polarity. We speculate that RomY *in vivo*, as observed *in vitro*, boosts MglB GAP activity to a sufficiently high level to outcompete RomR/RomX GEF activity at the lagging pole.

We only observed the interactions between RomY and MglB and MglA-GTP after protein crosslinking, suggesting that the three complexes that RomY can engage in the formation of are low affinity and transient. Overproduction of MglB *in vivo* in the presence of WT levels of RomY caused an increase in RomY/MglB GAP activity. Similarly, overproduction of RomY in the presence of WT levels of MglB caused an increase in RomY/MglB GAP activity. Altogether, these observations support that the rate-limiting step for RomY/MglB GAP activity in WT is the formation of the RomY/MglB complex rather than protein concentrations, corroborating that the RomY/MglB complex is a low affinity. *In vivo* RomY localizes significantly more asymmetrically to the lagging cell pole than MglB. Building on these observations, we suggest that RomY localizes highly asymmetrically to the lagging cell pole due to the high concentration of MglB at this pole. By contrast, RomY essentially does not localize to the leading pole because the concentration of MglB would be too low at this pole to support MglB/RomY complex formation. In principle, RomY could be recruited to the leading pole by MglA-GTP; however, we observed that MglA-GTP does not appear to play a role in the polar recruitment of RomY.

The RomR/RomX GEF and MglB GAP are both arranged intracellularly with a high concentration at the lagging pole and a low concentration at the leading pole. Nevertheless, GEF activity dominates at the leading and GAP activity at the lagging cell pole. It has been argued that this localization pattern is ideal to allow stable and switchable polarity and reflects a trade-off between maintaining stable polarity with unidirectional motility between reversals and sensitivity to Frz signaling with an inversion of polarity and cellular reversals ^36^. However, the “price” that cells pay for this design is the need for a mechanism to separate the GEF and GAP activities spatially. It has remained enigmatic how the spatial regulation of the GEF and GAP is brought about. The data presented here suggest that RomY is an elegant solution to this problem. Specifically, because the RomY/MglB complex is low-affinity, it is formed at the lagging pole with the high MglB concentration but not at the leading pole with the low MglB concentration. In this way, MglB GAP activity is precisely and only stimulated at the lagging pole.

Based on the data reported here, we suggest a revised model for the regulation of front-rear polarity in *M. xanthus*. In this model, MglA is activated and recruited to the leading pole by the RomR/RomX GEF complex; MglB at this pole is not active because RomY is absent. At the lagging pole, MglA-GTP is inactivated (i.e. MglA GTPase activity is activated) by the RomY/MglB GAP complex that specifically forms at this pole and outcompetes RomR/RomX activity (Fig. 6). Thus, the key to the spatially restricted activity of RomY/MglB is the low affinity of RomY for MglB.

**Figure 6.**
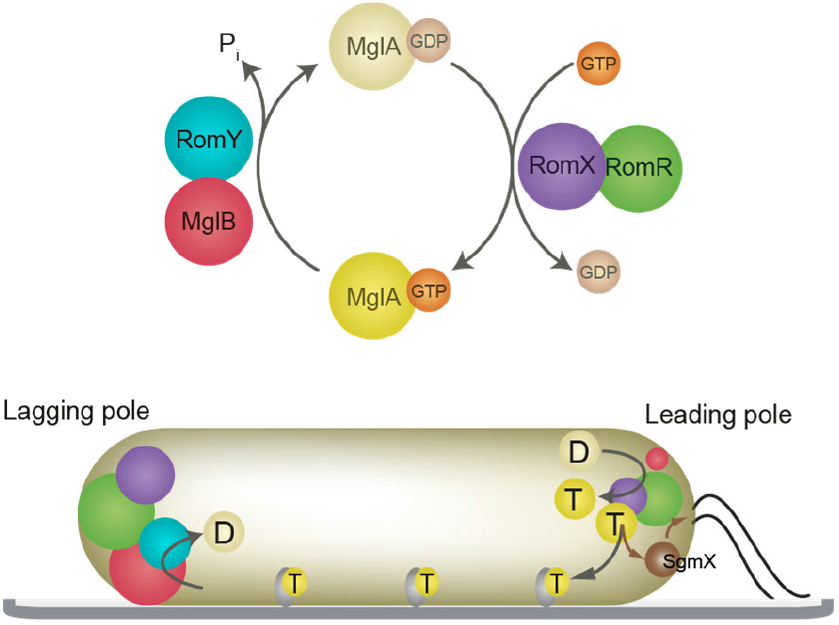
Model for front–rear polarity in *M. xanthus*. Upper panel, MglA GTPase cycle. The MglB/RomY complex is shown to indicate that both proteins interact with MglA-GTP and the RomR/RomX complex to indicate that only RomX interacts with MglA. Lower panel, localization of MglA-GTP, MglB, RomY, RomR, RomX and SgmX in a cell with T4P at the leading pole. Color code as in the upper panel, except that yellow circles labelled D and T represent MglA-GDP and MglA-GTP, respectively. SgmX is in brown and with the brown arrows indicating its recruitment by MglA-GTP and stimulation of T4P formation. The dark grey arrow indicates stimulation of assembly of the Agl/Glt complexes (light grey) and the incorporation of MglA-GTP into these complexes. Circle sizes indicate the amount of protein at a pole.

In eukaryotes, Rho GTPases are key regulators of motility and polarity and their activity is spatially confined to distinct intracellular locations. In some cases, this confinement has been shown to rely on spatially separated GEF and GAP activities. For instance, in *Drosophila* epithelial cells, Cdc42 colocalizes with its cognate GEFs at the apical membrane while the cognate GAP is at the lateral membrane and assists in restricting Cdc42 activity to the apical membrane ^44^. Thus, the design principles underlying polarity are overall similar in these systems and *M. xanthus*. However, in *M. xanthus*, polarity can be inverted, while it is stably maintained in epithelial cells. As mentioned, it has been suggested that the special arrangement with the RomR/RomX GEF in a “waiting position” at the lagging pole is key to this switchability ^36^. Because Ras-like GTPases are also involved in regulating dynamic polarity in eukaryotes ^5,8^, we speculate that polarity systems with a design similar to the *M. xanthus* system may underlie the regulation of dynamic polarity in eukaryotic cells.

Small Ras-like GTPases and Roadblock proteins are present in all three domains of life, and it has been suggested that they were present in the last universal common ancestor ^37,45–47^. Interestingly, proteins containing a Roadblock domain or the structurally related Longin domain, which might have evolved from the Roadblock domain ^48^, often form heteromeric complexes with GEF or GAP activity. For instance, the Ragulator complex has Rag GEF activity ^49–52^, the GATOR1 and FLCN/FNIP complexes Rag GAP activity ^53–55^, and the Mon1-Ccz1 and TRAPP-II complexes have Rab7 GEF ^56^ and Rab1 GEF ^57^ activity, respectively. Thus, the finding that MglB functions in a complex with RomY follows this theme and add the MglB/RomY complex to the list of heteromeric Roadblock domain-containing complexes important for regulating small GTPases. These observations also support the idea that cognate GTPase/Roadblock pairs could represent minimal, ancestral pairs of a GTPase and its regulator. During evolution, Roadblock domain-containing proteins would then have become incorporated into more complex GEFs and GAPs to regulate GTPase activity.

## Methods

### Cell growth and construction of strains

DK1622 was used as the WT *M. xanthus* strain and all strains are derivatives of DK1622. *M. xanthus* strains used are listed in Supplementary Table 2. Plasmids are listed in Supplementary Table 3. In-frame deletions were generated as described ^58^. *M. xanthus* was grown at 32°C in 1% casitone (CTT) broth ^59^ or on 1.5% agar supplemented with 1% CTT and kanamycin (50μg/ml) or oxytetracycline (10μg/ml) if appropriate. Plasmids were integrated by site specific recombination into the Mx8 *attB* site or by homologous recombination at the native site. All in-frame deletions and plasmid integrations were verified by PCR. Primers used are listed in Supplementary Table 4. Plasmids were propagated in *Escherichia coli* TOP10 (F^-^, *mcrA*, Δ(*mrr-hsd*RMS-*mcr*BC), φ80*lac*ZΔM15, Δ*lac*X74, *deo*R, *rec*A1, *ara*D139, Δ(*ara-leu*)7679, *gal*U, *gal*K, *rps*L, *endA*1, *nup*G) unless otherwise stated. *E. coli* cells were grown in LB or on plates containing LB supplemented with 1.5% agar at 37 °C with added antibiotics if appropriate ^60^. All DNA fragments generated by PCR were verified by sequencing.

### Motility assays and determination of reversal frequency

Population-based motility assays were done as described ^38^. Briefly, *M. xanthus* cells from exponentially growing cultures were harvested at 4000× *g* for 10 min at room temperature (RT) and resuspended in 1% CTT to a calculated density of 7×10^9^ cells ml^-1^. 5μL aliquots of cell suspensions were placed on 0.5% agar plates supplemented with 0.5% CTT for T4P-dependent motility and 1.5% agar plates supplemented with 0.5% CTT for gliding motility and incubated at 32°C. After 24h, colony edges were visualized using a Leica M205FA stereomicroscope and imaged using a Hamamatsu ORCA-flash V2 Digital CMOS camera (Hamamatsu Photonics). For higher magnifications of cells at colony edges on 1.5% agar, cells were visualized using a Leica DMi8 inverted microscope and imaged with a Leica DFC9000 GT camera. Individual cells were tracked as described ^29^. Briefly, for T4P-dependent motility, 5μL of exponentially growing cultures were spotted into a 24-well polystyrene plate (Falcon). After 10min at RT, cells were covered with 500μL of 1% methylcellulose in MMC buffer (10mM MOPS (3-(*N*-morpholino)propanesulfonic acid) pH 7.6, 4mM MgSO_4_, 2mM CaCl_2_), and incubated at RT for 30min. Subsequently, cells were visualized for 10min at 20sec intervals at RT using a Leica DMi8 inverted microscope and a Leica DFC9000 GT camera. Individual cells were tracked using Metamorph 7.5 (Molecular Devices) and ImageJ 1.52b ^61^ and then the speed of individual cells per 20sec interval as well as the number of reversals per cell per 10min calculated. For gliding, 5μL of exponentially growing cultures were placed on 1.5% agar plates supplemented with 0.5% CTT, covered by a cover slide and incubated at 32°C. After 4 to 6h, cells were observed for 15min at 30sec intervals at RT as described above and then the fraction of moving cells, speed per 30sec interval as well as the number of reversals per 15min calculated.

For experiment with vanillate, cells were diluted to the same optical density (OD) at 550nm of 0.2, grown for 30min at 32°C in suspension culture, and then vanillate was added to a final concentration of 500μM. Subsequently, cells were grown 3h at 32°C before cells were spotted into a 24-well polystyrene plate (Falcon). After 10min at RT, cells were covered with 500μL of 1% methylcellulose in MMC buffer supplemented with 500μM vanillate, and incubated at RT for 30min. Subsequently, cells were visualized for 10min at 20sec intervals at RT as described. Control cultures without vanillate were treated similarly.

### Fluorescence microscopy

Epifluorescence microscopy was done as described ^29^. Briefly, *M. xanthus* cells were placed on a thin 1.5% agar pad buffered with TPM buffer (10mM Tris-HCl pH 8.0, 1mM potassium phosphate buffer pH 7.6, 8mM MgSO_4_) on a glass slide and immediately covered with a coverslip. After 30min at 32°C, cells were visualized using a Leica DMi8 microscope and imaged with Hamamatsu ORCA-flash V2 Digital CMOS camera. Cells in phase contrast images were automatically detected using Oufti ^62^. Fluorescence signals in segmented cells were identified and analyzed using a custom-made Matlab v2016b (MathWorks) script ^29^. Briefly, polar clusters were identified when they had an average fluorescence two STDEV above the average cytoplasmic fluorescence and a size of three or more pixels. For each cell with polar clusters, an asymmetry index (ω) was calculated as

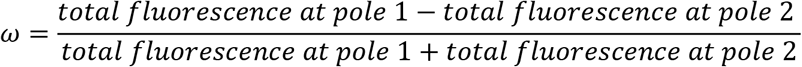

By definition, pole 1 is the pole with the highest fluorescence. ω varies between 0 (bipolar symmetric localization) and 1 (unipolar localization). The localization patterns were binned from the ω values as follows: unipolar (ω > 0.9), bipolar asymmetric (0.9 > ω > 0.2) and bipolar symmetric (ω<0.2). Diffuse localization was determined when no polar signal was detected.

For time-lapse epifluorescence microscopy, cells were prepared as described. Time-lapse recordings were made for 15min with images recorded every 30sec. Data were processed with Metamorph 7.5 and ImageJ 1.52b. Cells in phase contrast images were automatically detected using Oufti. Fluorescence signals in segmented cells were identified and analyzed using a custom-made Matlab script. Briefly, polar clusters were identified when they had an average fluorescence two STDEV above the average cytoplasmic fluorescence, an average fluorescence two-fold higher than the average the cytoplasmic fluorescence, and a size of three or more pixels. A custom-made Matlab script was used to track cells, detect reversals, leading and lagging cell poles, and to plot the data.

### Immunoblot analysis

Immunoblots were done as described ^60^. Rabbit polyclonal antibodies α-MglA ^23^, α-MglB ^23^, α-PilC ^20^ and α-RomY antibodies were used together with goat anti-rabbit immunoglobulin G conjugated with horseradish peroxidase (Sigma) as secondary antibody. Monoclonal mouse anti-polyHistidine antibodies conjugated with peroxidase (Sigma) were used to detect His_6_ tagged proteins. To generate rabbit, polyclonal α-RomY antibodies, purified His_6_-RomY was used to immunize rabbit as described ^60^. Blots were developed by using Luminata Crescendo Western HRP Substrate (Millipore) and visualized using a LAS-4000 luminescent image analyzer (Fujifilm).

### Protein purification

All proteins were expressed in *E. coli* Rosetta 2(DE3) (F^-^ *ompThsdS_B_*(r_B_^-^ m_B_^-^) *gal dcm* (DE3 pRARE2) at 18°C or 37°C. To purify His_6_-tagged proteins, Ni-NTA affinity purification was used. Briefly, cells were washed in buffer A (50mM Tris pH 7.5, 150mM NaCl, 10mM imidazole, 5%glycerol, 5mM MgCl_2_) and resuspended in lysis buffer A (50 ml of wash buffer A supplemented with 1mM DTT, 100μg mL^-1^ phenylmethylsulfonyl fluoride (PMSF), 10U mL^-1^ DNase 1 and protease inhibitors – Complete Protease Inhibitor Cocktail Tablet (Roche)). Cells were lysed by sonication, cell debris removed by centrifugation (48000× *g*, 4°C, 30min), and cell lysate filtered through 0.45 μm Polysulfone filter (Filtropur S 0.45, Sarstedt). The cleared cell lysate was loaded onto a 5mL HiTrap Chelating HP column (GE Healthcare) preloaded with NiSO_4_ as described by the manufacturer and equilibrated in buffer A. The column was washed with 20 column volumes of buffer A supplemented with 20mM imidazole. Proteins were eluted with buffer A using a linear imidazole gradient from 20-500mM. Fractions containing purified MglA-His_6_ or His_6_-MglB proteins were combined and loaded onto a HiLoad 16/600 Superdex 75 pg (GE Healthcare) gel filtration column that was equilibrated with buffer A without imidazole for use in GTPase assays or buffer C (20mM HEPES (4-(2-hydroxyethyl)-1-piperazineethanesulfonic acid) pH 8.0, 150mM NaCl, 5mM MgCl_2_) for use in pull-down experiments. Fractions containing His_6_-MalE were combined and loaded on a HiLoad 16/600 Superdex 200 pg (GE Healthcare) column equilibrated with buffer C. Fractions containing His_6_-tagged proteins were pooled, frozen in liquid nitrogen and stored at −80°C.

To purify Strep-RomY and Strep-RomY^N^, biotin affinity purification was used. Briefly, cells were washed in buffer D (100mM Tris pH 8.0, 150mM NaCl, 1mM EDTA, 1mM DTT) and resuspended in lysis buffer D (50 ml of buffer D supplemented with 100μg mL^-1^ PMSF, 10U mL^-1^ DNase 1 and protease inhibitors – Complete Protease Inhibitor Cocktail Tablet (Roche)). Cells were lysed and cleared lysate prepared as described and loaded onto a 5 mL Strep-Trap HP column (GE Healthcare), equilibrated with buffer D. The column was washed with 20 column volumes of buffer D. Protein was eluted with buffer E (150mM Tris pH 8.0, 150mM NaCl, 1mM EDTA, 2.5mM Desthiobiotin). Elution fractions containing Strep-RomY or StrepRomY^N^ were loaded onto a a HiLoad 16/600 Superdex 200 pg (GE Healthcare) gel filtration column that was equilibrated with buffer A without imidazole for use in GTPase assays or buffer C for use in pull-down experiments. Fractions with Strep-RomY or StrepRomY^N^ were pooled, frozen in liquid nitrogen and stored at −80°C.

### GTPase assays

GTP-hydrolysis by MglA-His_6_ was measured using a continuous, regenerative coupled GTPase assay ^63^ or by measuring released inorganic phosphate (P_i_) after GTP hydrolysis using a malachite green assay ^64^. The continuous, regenerative coupled GTPase assay was performed in buffer F (50mM Tris pH 7.5, 150mM NaCl, 5% glycerol, 1mM DTT, 7.5mM MgCl_2_) supplemented with 495μM NADH (Sigma), 2mM phosphoenolpyruvate (Sigma), 18-30U mL^-1^ pyruvate kinase (Sigma) and 27-42 U mL^-1^ lactate dehydrogenase (Sigma). MglA-His_6_ (final concentration: 10μM) was pre-loaded with GTP (final concentration: 3.3mM) for 30min at RT in buffer F. In parallel, His_6_-MglB, Strep-RomY, Strep-RomY^N^ or equimolar amount of His_6_-MglB and Strep-RomY/Strep-RomY^N^ (final concentrations of all proteins: 8.6 μM) were preincubated for 10min at RT in buffer F. Reactions were started in a 96-well plate (Greiner Bio-One) by adding His_6_-MglB and/or Strep-RomY/Strep-RomY^N^ to the MglA/GTP mixture. Final concentrations in these reactions: MglA-His_6_: 3μM, His_6_-MglB: 6μM, Strep-RomY/Strep-RomY^N^: 6μM, GTP: 1mM. Absorption was measured at 340nm for 60min at 37°C with an Infinite M200 Pro plate-reader (Tecan) and the amount of hydrolyzed GTP per h per molecule of MglA-His_6_ calculated. For each reaction, background subtracted GTPase activity was calculated as the mean of three technical replicates. In the malachite green assay, released P_i_ during GTP hydrolysis was measured in buffer F. Proteins were used in concentrations and preincubated as described. GTPase reactions were performed in 96-well plates (Greiner Bio-One) at 37°C and started by adding His_6_-MglB and/or Strep-RomY/Strep-RomY^N^ to the MglA/GTP mixture. Final concentrations as described. After 1h, reactions were stopped and the colour developed according to the manufacturer’s manual (BioLegend) and absorption at 590nm measured using an Infinite M200 Pro plate-reader (Tecan). Subsequently, released P_i_ was calculated from a standard curve, and the amount of released P_i_ per h per MglA-His_6_ molecule calculated.

### Pull-down experiments

In all experiments involving MglA-His_6_, MglA-His_6_ was preloaded with GTP or GDP (44.4μM protein, 22.2mM GTP/GDP) for 30min at RT in buffer C. Subsequently, equimolar amounts of Strep-RomY and MglA-His_6_, His_6_-MglB or His_6_-MalE were incubated for 30min RT in buffer C. Final concentrations: MglA-His_6_, His_6_-MglB, His_6_-MalE, Strep-RomY: 20μM, GTP/GDP 10mM. Where indicated, DSP was added to a final concentration of 200μM for 5min at RT. Next, all reactions were quenched with Tris pH 7.6 added to a final concentration of 100mM and incubated for 15min at RT. Subsequently, 20μl of Strep-Tactin coated magnetic beads (MagStrep ‘type3’ XT beads (IBA-Lifesciences)) previously equilibrated with buffer C were added and samples incubated for 30min RT. The beads were washed 10 times with 1mL buffer C. For experiments with GTP or GDP, buffer C was supplemented with 5mM GTP/GDP. Proteins were eluted with 100μL elution buffer (100 mM Tris pH 8.0, 150mM NaCl, 1mM EDTA, 50mM biotin). Samples were prepared in SDS-PAGE loading buffer (60mM Tris pH 6.8, 2% SDS, 10% glycerol, 0.005% bromophenol blue, 5 mM EDTA) with or without 100mM DTT (final concentration) as indicated. In all SDS–PAGE experiments, equivalent volumes of loading and wash fractions and two-fold more of the elution fraction were loaded and gels stained with Coomassie Brilliant Blue and subsequently analyzed by immunoblotting.

### AlphaFold structural models

AlphaFold and AlphaFold-multimer structure prediction was done with the ColabFold pipeline ^41–43^. ColabFold was executed with default settings where multiple sequence alignments were generated with MMseqs2 ^65^ and HHsearch ^66^. The ColabFold pipeline generates five model ranks. Predicted Local Distance Difference Test (pLDDT) and alignment error (pAE) graphs were generated for each rank with custom Matlab script. Models of the highest confidence based on combined pLDDT and pAE values were used for further investigation and presentation. Structural alignments and images were generated in Pymol (The PyMOL Molecular Graphics System, Version 1.2r3pre, Schrödinger, LLC). For all models, sequences of full-length proteins were used.

### Bioinformatics

Sequence alignments were done using MUSCLE ^67^ with default parameters in MEGA7 ^68^ and alignments were visualized with GeneDoc ^69^. Protein domains were identified using SMART ^70^. % similarity/identity between protein homologs were calculated using EMBOSS Needle software (pairwise sequence alignment) ^71^.

### Statistics

Statistics were performed using a two-tailed Student’s *t*-test for samples with unequal variances.

### Data availability

The authors declare that all data supporting this study are available within the article and its Supplementary Information file. The source data underlying Fig. 1b, c, d, e, 2a, b, 3c, 4a, b, 5a, c, d and Supplementary Fig. 2, 3a, b, c, 4a (pLLDT and pAE for selected rank model of RomY, MglA:(MglB)_2_, MglA:RomY, (MglB)_2_:RomY and MglA:(MglB)_2_:RomY), 5b, 6, 7b, c, d, e, f are provided as a Source Data file.

### Code availability

The Matlab scripts used in this study are available from the corresponding author upon request.

## Acknowledgements

We gratefully acknowledge the help of Marco Herfurth and Dr. Dorota Skotnicka for many helpful discussions and of Dr. Kristin Wuichet and Dr. Daniela Keilberg in the identification and initial characterization of RomY. This work was funded by the Deutsche Forschungsgemeinschaft (project no. 269423233) within the framework of the Transregio 174 ‘Spatiotemporal dynamics of bacterial cells’ (to L.S.-A.) as well as by the Max Planck Society.

## Funding

This work was supported by the Max Planck Society (to LSA).

## Author contributions

D.S, L.A.M.C and L.S.-A. conceptualized the study.

D.S. and L.A.M.C. conducted the experimental work.

D.S. and L.A.M.C. analyzed experimental data.

D.S. and L.S.-A. wrote the original draft of the manuscript.

D.S., L.A.M.C., and L.S.-A. reviewed and edited the manuscript.

L.S.-A. provided supervision.

L.S.-A. acquired funding.

## Declaration of Interests

The authors declare no competing interests.

## Notes

### Competing Interest Statement

The authors have declared no competing interest.

## References

1 Rafelski, S. & Marshall, W. Building the cell: design principles of cellular architecture. Nat Rev Mol Cell Biol 9, 593–602 (2008).

2 Treuner-Lange, A. & Søgaard-Andersen, L. Regulation of cell polarity in bacteria. J. Cell Biol. 206, 7–17 (2014).

3 Surovtsev, I. V. & Jacobs-Wagner, C. Subcellular organization: A critical feature of bacterial cell replication. Cell 172, 1271–1293 (2018).

4 Chiou, J.-G., Balasubramanian, M. K. & Lew, D. J. Cell polarity in yeast. Ann. Rev. Cell Dev. Biol. 33, 77–101 (2017).

5 Etienne-Manneville, S. & Hall, A. Rho GTPases in cell biology. Nature 420, 629–635 (2002).

6 Wu, C.-F. & Lew, D. J. Beyond symmetry-breaking: competition and negative feedback in GTPase regulation. Trends Cell Biol 23, 476–483 (2013).

7 Schumacher, D. & Søgaard-Andersen, L. Regulation of cell polarity in motility and cell division in Myxococcus xanthus. Annu. Rev. Microbiol. 71, 61–78 (2017).

8 Ridley, A. J. et al. Cell migration: Integrating signals from front to back. Science 302, 1704–1709 (2003).

9 Spang, A. et al. Complex archaea that bridge the gap between prokaryotes and eukaryotes. Nature 521, 173–179 (2015).

10 Vetter, I. R. & Wittinghofer, A. The guanine nucleotide-binding switch in three dimensions. Science 294, 1299–1304 (2001).

11 Cherfils, J. & Zeghouf, M. Regulation of small GTPases by GEFs, GAPs, and GDIs. Physiol. Rev. 93, 269–309 (2013).

12 Bos, J. L., Rehmann, H. & Wittinghofer, A. GEFs and GAPs: Critical elements in the control of small G proteins. Cell 129, 865–877 (2007).

13 Zhang, Y., Ducret, A., Shaevitz, J. & Mignot, T. From individual cell motility to collective behaviors: insights from a prokaryote, *Myxococcus xanthus*. FEMS Microbiol. Rev. 36, 149–164 (2012).

14 Blackhart, B. D. & Zusman, D. R. “Frizzy” genes of *Myxococcus xanthus* are involved in control of frequency of reversal of gliding motility. Proc. Natl. Acad. Sci. USA 82, 8771–8774 (1985).

15 Mignot, T., Shaevitz, J. W., Hartzell, P. L. & Zusman, D. R. Evidence that focal adhesion complexes power bacterial gliding motility. Science 315, 853–856 (2007).

16 Treuner-Lange, T. et al. The small G-protein MglA connects to the MreB actin cytoskeleton at bacterial focal adhesions. J. Cell Biol. 210, 243–256 (2015).

17 Sun, H., Zusman, D. R. & Shi, W. Type IV pilus of *Myxococcus xanthus* is a motility apparatus controlled by the *frz* chemosensory system. Curr. Biol. 10, 1143–1146 (2000).

18 Merz, A. J., So, M. & Sheetz, M. P. Pilus retraction powers bacterial twitching motility. Nature 407, 98–102 (2000).

19 Skerker, J. M. & Berg, H. C. Direct observation of extension and retraction of type IV pili. Proc. Natl. Acad. Sci. USA 98, 6901–6904 (2001).

20 Bulyha, I. et al. Regulation of the type IV pili molecular machine by dynamic localization of two motor proteins. Mol. Microbiol. 74, 691–706 (2009).

21 Hodgkin, J. & Kaiser, D. Genetics of gliding motility in *Myxococcus xanthus* (Myxobacterales): Two gene systems control movement. Mol Gen Genet 171, 177–191 (1979).

22 Hartzell, P. & Kaiser, D. Function of MglA, a 22-kilodalton protein essential for gliding in *Myxococcus xanthus*. J Bacteriol 173, 7615–7624 (1991).

23 Leonardy, S. et al. Regulation of dynamic polarity switching in bacteria by a Ras-like G-protein and its cognate GAP. EMBO J. 29, 2276–2289 (2010).

24 Zhang, Y., Franco, M., Ducret, A. & Mignot, T. A bacterial Ras-like small GTP-binding protein and its cognate GAP establish a dynamic spatial polarity axis to control directed motility. PLOS Biol 8, e1000430 (2010).

25 Mauriello, E. M. F. et al. Bacterial motility complexes require the actin-like protein, MreB and the Ras homologue, MglA. EMBO J 29, 315–326 (2010).

26 Yang, R. et al. AglZ is a filament-forming coiled-coil protein required for adventurous motility of *Myxococcus xanthus*. J. Bacteriol. 186, 6168–6178 (2004).

27 Potapova, A., Carreira, L. A. M. & Søgaard-Andersen, L. The small GTPase MglA together with the TPR domain protein SgmX stimulates type IV pili formation in *M. xanthus*. Proc. Natl. Acad Sci. USA 117, 23859–23868 (2020).

28 Mercier, R. et al. The polar Ras-like GTPase MglA activates type IV pilus via SgmX to enable twitching motility in *Myxococcus xanthus*. Proc. Natl. Acad Sci. USA 117, 28366–28373 (2020).

29 Szadkowski, D. et al. Spatial control of the GTPase MglA by localized RomR/RomX GEF and MglB GAP activities enables *Myxococcus xanthus* motility. Nat. Microbiol. 4, 1344–1355 (2019).

30 Baranwal, J. et al. Allosteric regulation of a prokaryotic small Ras-like GTPase contributes to cell polarity oscillations in bacterial motility. PLOS Biol. 17, e3000459 (2019).

31 Miertzschke, M. et al. Structural analysis of the Ras-like G protein MglA and its cognate GAP MglB and implications for bacterial polarity. EMBO J. 30, 4185–4197 (2011).

32 Galicia, C. et al. MglA functions as a three-state GTPase to control movement reversals of *Myxococcus xanthus*. Nat. Comm. 10, 5300 (2019).

33 Keilberg, D., Wuichet, K., Drescher, F. & Søgaard-Andersen, L. A response regulator interfaces between the Frz chemosensory system and the MglA/MglB GTPase/GAP module to regulate polarity in *Myxococcus xanthus*. PLOS Genet. 8, e1002951 (2012).

34 Leonardy, S., Freymark, G., Hebener, S., Ellehauge, E. & Søgaard-Andersen, L. Coupling of protein localization and cell movements by a dynamically localized response regulator in *Myxococcus xanthus*. EMBO J. 26, 4433–4444 (2007).

35 Zhang, Y., Guzzo, M., Ducret, A., Li, Y.-Z. & Mignot, T. A dynamic response regulator protein modulates G-protein–dependent polarity in the bacterium *Myxococcus xanthus*. PLOS Genet 8, e1002872 (2012).

36 Carreira, L. A. M., Tostevin, F., Gerland, U. & Søgaard-Andersen, L. Protein-protein interaction network controlling establishment and maintenance of switchable cell polarity. PLOS Genetics 16, e1008877 (2020).

37 Wuichet, K. & Søgaard-Andersen, L. Evolution and diversity of the Ras superfamily of small GTPases in prokaryotes. Genome Biol Evol 7, 57–70 (2015).

38 Shi, W. & Zusman, D. R. The two motility systems of *Myxococcus xanthus* show different selective advantages on various surfaces. Proc. Natl. Acad. Sci. USA 90, 3378–3382 (1993).

39 Blackhart, B. D. & Zusman, D. R. Analysis of the products of the *Myxococcus xanthus frz* genes. J. Bacteriol. 166, 673–678 (1986).

40 Guzzo, M. et al. Evolution and design governing signal precision and amplification in a bacterial chemosensory pathway. PLOS Genet. 11, e1005460 (2015).

41 Jumper, J. et al. Highly accurate protein structure prediction with AlphaFold. Nature 596, 583–589 (2021).

42 Mirdita, M., Ovchinnikov, S. & Steinegger, M. ColabFold - Making protein folding accessible to all. bioRxiv, 2021.2008.2015.456425 (2021).

43 Evans, R. et al. Protein complex prediction with AlphaFold-Multimer. bioRxiv, 2021.2010.2004.463034 (2021).

44 Fic, W. et al. RhoGAP19D inhibits Cdc42 laterally to control epithelial cell shape and prevent invasion. J. Cell Biol. 220 (2021).

45 Koonin, E. V. & Aravind, L. Dynein light chains of the Roadblock/LC7 group belong to an ancient protein superfamily implicated in NTPase regulation. Curr. Biol. 10, R774–R776 (2000).

46 Klinger, C. M., Spang, A., Dacks, J. B. & Ettema, T. J. G. Tracing the Archaeal origins of eukaryotic membrane-trafficking system building blocks. Mol. Biol. Evol. 33, 1528–1541 (2016).

47 Liu, Y. et al. Expanded diversity of Asgard archaea and their relationships with eukaryotes. Nature 593, 553–557 (2021).

48 Levine, T. P. et al. Discovery of new Longin and Roadblock domains that form platforms for small GTPases in Ragulator and TRAPP-II. Small GTPases 4, 62–69 (2013).

49 Zhang, T. et al. Structural basis for Ragulator functioning as a scaffold in membrane-anchoring of Rag GTPases and mTORC1. Nat. Comm. 8, 1394 (2017).

50 Bar-Peled, L., Schweitzer, L. D., Zoncu, R. & Sabatini, D. M. Ragulator is a GEF for the Rag GTPases that signal amino acid levels to mTORC1. Cell 150, 1196–1208 (2012).

51 de Araujo, M. E. G. et al. Crystal structure of the human lysosomal mTORC1 scaffold complex and its impact on signaling. Science 358, 377–381 (2017).

52 Su, M.-Y. et al. Hybrid structure of the RagA/C-Ragulator mTORC1 activation complex. Mol. Cell 68, 835–846.e833 (2017).

53 Lawrence, R. E. et al. Structural mechanism of a Rag GTPase activation checkpoint by the lysosomal folliculin complex. Science 366, 971–977 (2019).

54 Shen, K. et al. Architecture of the human GATOR1 and GATOR1–Rag GTPases complexes. Nature 556, 64–69 (2018).

55 Shen, K. et al. Cryo-EM structure of the human FLCN-FNIP2-Rag-Ragulator complex. Cell 179, 1319–1329.e1318 (2019).

56 Nordmann, M. et al. The Mon1-Ccz1 complex is the GEF of the late endosomal Rab7 homolog Ypt7. Curr. Biol. 20, 1654–1659 (2010).

57 Yip, C. K., Berscheminski, J. & Walz, T. Molecular architecture of the TRAPPII complex and implications for vesicle tethering. Nat. Struct. Mol. Biol. 17, 1298–1304 (2010).

58 Shi, X. et al. Bioinformatics and experimental analysis of proteins of two-component systems in *Myxococcus xanthus*. J. Bacteriol. 190, 613–624 (2008).

59 Hodgkin, J. & Kaiser, D. Cell-to-cell stimulation of movement in nonmotile mutants of *Myxococcus*. Proc. Natl. Acad. Sci. USA 74, 2938–2942 (1977).

60 Sambrook, J. & Russell, D. W. Molecular Cloning: A Laboratory Manual. 3rd edn, (Cold Spring Harbor Laboratory Press), (2001).

61 Schneider, C. A., Rasband, W. S. & Eliceiri, K. W. NIH Image to ImageJ: 25 years of image analysis. Nat. Methods 9, 671–675 (2012).

62 Paintdakhi, A. et al. Oufti: an integrated software package for high-accuracy, high-throughput quantitative microscopy analysis. Mol. Microbiol. 99, 767–777 (2016).

63 Ingerman, E. & Nunnari, J. A continuous, regenerative coupled GTPase assay for dynamin-related proteins. Methods Enzymol 404, 611–619 (2005).

64 Lanzetta, P. A., Alvarez, L. J., Reinach, P. S. & Candia, O. A. An improved assay for nanomole amounts of inorganic phosphate. Anal. Biochem. 100, 95–97 (1979).

65 Mirdita, M., Steinegger, M. & Söding, J. MMseqs2 desktop and local web server app for fast, interactive sequence searches. Bioinformatics 35, 2856–2858 (2019).

66 Steinegger, M. et al. HH-suite3 for fast remote homology detection and deep protein annotation. BMC Bioinformatics 20, 473 (2019).

67 Thompson, J. D., Higgins, D. G. & Gibson, T. J. CLUSTAL W: improving the sensitivity of progressive multiple sequence alignment through sequence weighting, positions-specific gap penalties and weight matrix choice. Nucl. Acids Res. 22, 4673–4680 (1994).

68 Kumar, S., Stecher, G. & Tamura, K. MEGA7: Molecular evolutionary genetics analysis Version 7.0 for bigger datasets. Mol Biol Evol 33, 1870–1874 (2016).

69 Nicholas, K. B., Nicholas Jr., H. B. & Deerfield II, D. W. Genedoc: Analysis and visualization of genetic variation. EMBNEW.NEWS 4, 14 (1997).

70 Letunic, I. & Bork, P. 20 years of the SMART protein domain annotation resource. Nucl. Acids Res. 46, D493–D496 (2017).

71 Li, W. et al. The EMBL-EBI bioinformatics web and programmatic tools framework. Nucl Acids Res 43, W580–W584 (2015).

