## Supplementary Information for "A bipartite, low-affinity roadblock domain-containing GAP complex regulates bacterial front-rear polarity"

35043 Marburg

Germany

**This file contains**

- Supplementary Figures 1-7
- Supplementary Tables 1-4
- Supplementary References

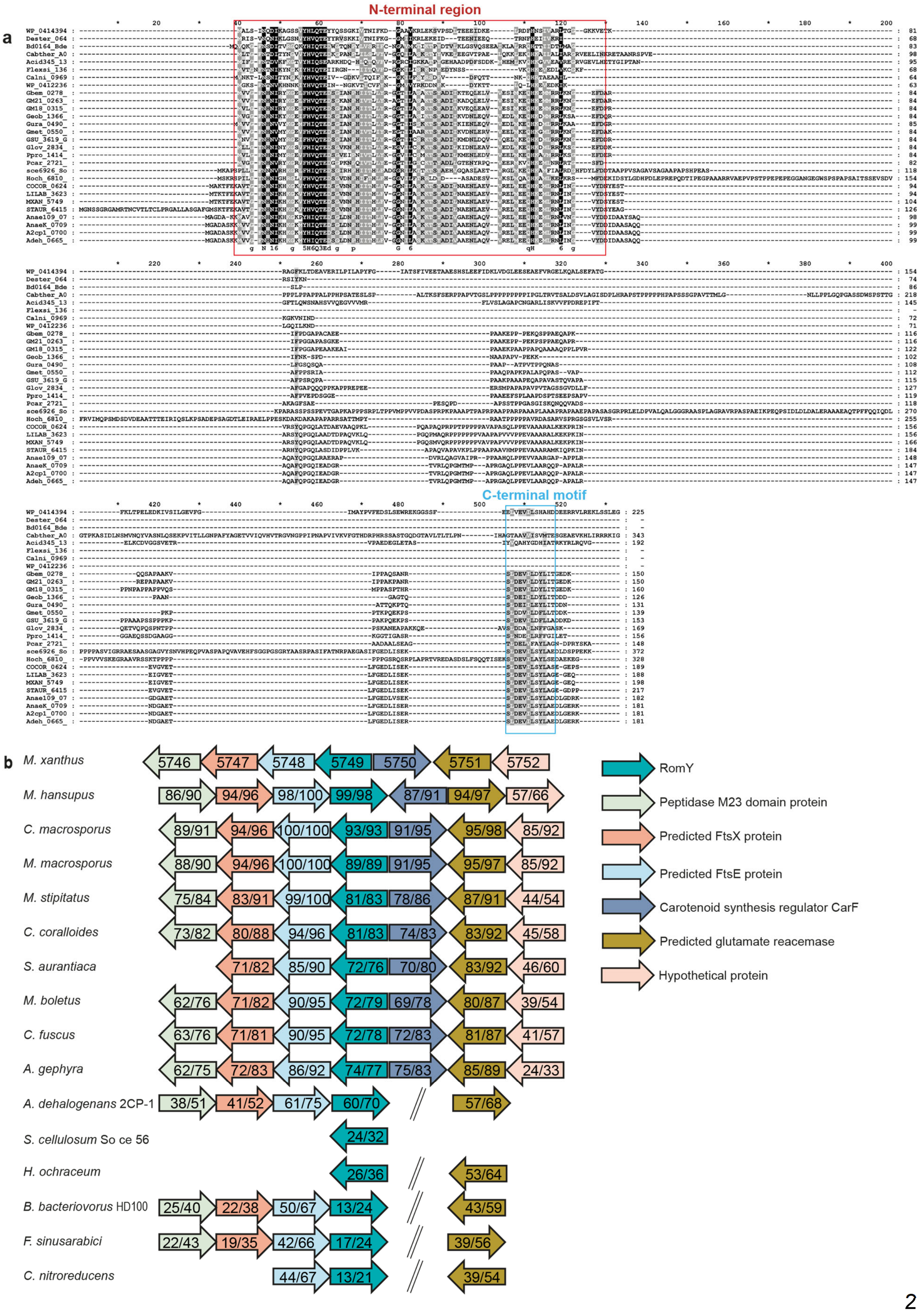
a
N-terminal region
C-terminal motif
b
M. xanthus
M. hansupus
C. macrosporus
M. macrosporus
M. stipitatus
C. coralloides
S. aurantiaca
M. boletus
C. fuscus
A. gephyra
A. dehalogenans 2CP-1
S. cellulosum So ce 56
H. ochraceum
B. bacteriovorus HD100
F. sinusarabici
C. nitroreducens
RomY
Peptidase M23 domain protein
Predicted FtsX protein
Predicted FtsE protein
Carotenoid synthesis regulator CarF
Predicted glutamate reacemase
Hypothetical protein

**Figure S1. The RomY protein and the *romY* locus**

**a.** Sequence alignment of RomY homologs. In red, the conserved N-terminal region, and light blue, the partially conserved C-terminal motif.

**b.** The *romY* locus is partially conserved. Transcription direction is indicated by the orientation of arrows with MXAN numbers indicated for the *romY* locus in *M. xanthus*. % similarity/identity between homologs from *M. xanthus* and other species is indicated by numbers in the arrows. For the proteins encoded by genes flanking *romY* in *M. xanthus*, domains were identified using SMART[1]. % similarity/identity between protein homologs were calculated using EMBOSS Needle software (pairwise sequence alignment) [2]. All listed species belong to the order Myxococcales except for *Bdellovibrio bacteriovorus* HD100 that belongs to the class Oligoflexia and *Flexistipes sinusarabica* and *Calditerrivibrio nitroreducens* that belong to the class Deferribacteres.

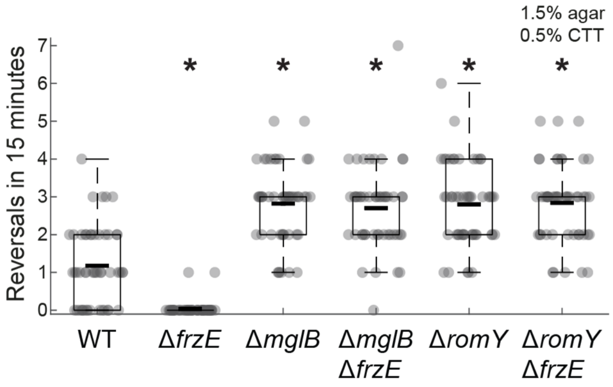


**Figure S2. Lack of RomY causes hyper-reversals independently of the Frz system**
Cells were incubated on 1.5% agar supplemented with 0.5% CTT to score gliding motility. Single data point are plotted in gray. Boxplots of reversals per cell in 15 min; boxes enclose 25$^{th}$ and 75$^{th}$ percentiles, cyan lines indicate the median, and whiskers the 10$^{th}$ and 90$^{th}$ percentiles. In all panels, * $P$< 0.01, two-sided Student's *t*-test.
Source data are provided in Source Data file.

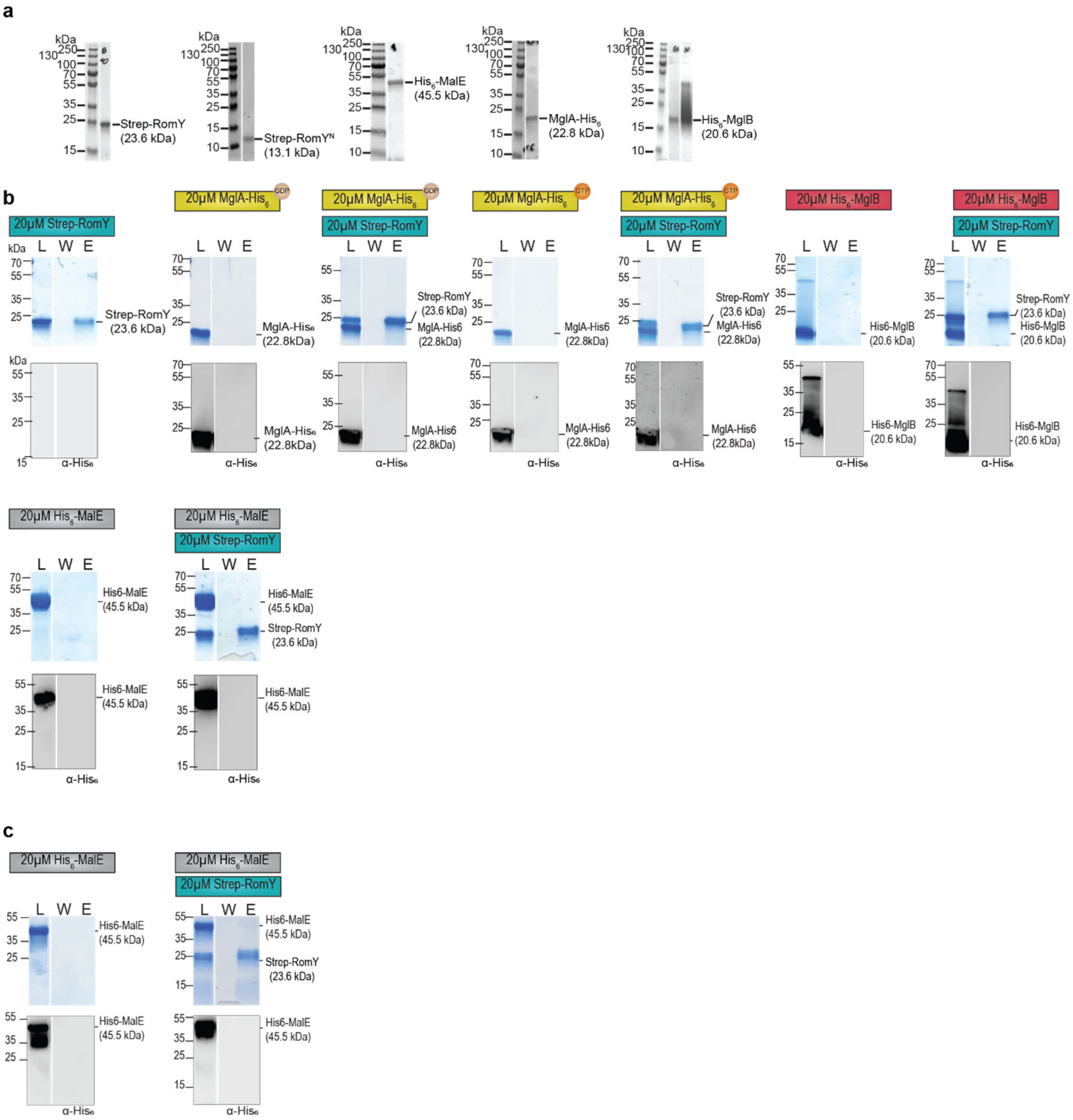


**Figure S3. RomY interaction with MglA-GTP and MglB is not detected in the absence of DSP crosss-linking**

**a.** SDS-PAGE analysis of purified proteins used in *in vitro* assays. ~5-20ng of the indicated purified proteins were separated by SDS-PAGE and gels stained with Coomassie Brilliant Blue. Calculated molecular weight of the different proteins is indicated. Molecular size markers are indicated on the left. Similar results were obtained in two independent experiments.

**b.** RomY interaction with MglA-GTP and MglB is not detected in the absence of DSP crosss-linking. Proteins were mixed with final concentrations and 10mM GTP/GDP as indicated in the schematics for 30min at RT, and proteins applied to Strep-Tactin coated magnetic beads.

Fractions before loading (L), the last wash (W) and after elution (E) were separated by SDS–PAGE, gels stained with Coomassie Brilliant Blue (upper panels) and subsequently probed with α-$His_6$ antibodies (lower panels). All samples were prepared with loading dye supplemented with 100mM DTT. For each combination, fractions were separated on the same gel. Gaps between lanes indicate lanes deleted for presentation purposes. Similar results were obtained in two independent experiments.

**c.** RomY does not interact with $His_6$-MalE after DSP cross-linking. Proteins were mixed with final concentrations as indicated in the schematics for 30min at RT, DSP added (final concentration 200μM, 5min, RT), DSP quenched, and proteins applied to Strep-Tactin coated magnetic beads. Fractions before loading (L), the last wash (W) and after elution (E) were separated by SDS–PAGE, gels stained with Coomassie Brilliant Blue (upper panels) and subsequently probed with α-$His_6$ antibodies (lower panels). All samples were treated with loading buffer containing 100mM DTT to break crosslinks before SDS-PAGE. For each combination, fractions were separated on the same gel. Gaps between lanes indicate lanes deleted for presentation purposes. Similar results were obtained in two independent experiments.

Source data for **a-c** are provided in Source Data file.

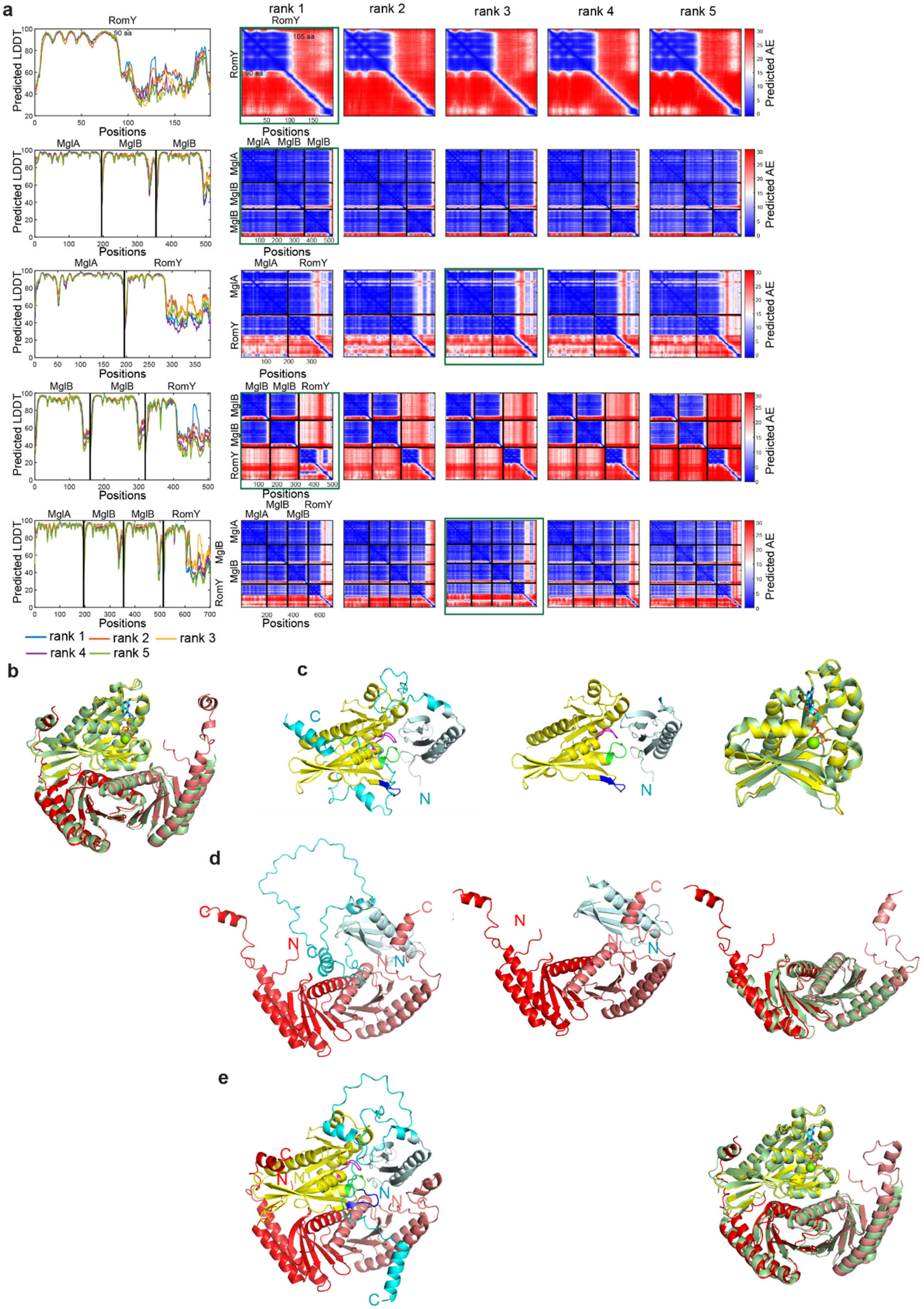

a
rank 1
rank 2
rank 3
rank 4
rank 5
RomY
MglA
MglB
Predicted LDDT
Positions
Predicted AE
90 aa
105 aa
rank 1 rank 2 rank 3
rank 4 rank 5
b
c
d
e
N
C

**Figure S4.** AlphaFold and AlpfaFold-Multimer models of RomY, MglA:RomY, $(MglB)_2$:RomY, MglA:$(MglB)_2$:RomY and MglA:$(MglB)_2$.

**a.** pLDDT and pAE plots of five models of the indicated protein and complexes. The model rank shown in Figure **3a,b** and **b-e** are indicated by a green box. AlphaFold was used for modeling RomY and AlphaFold-Multimer for modeling the complexes.

**b.** AlphaFold-Multimer model of MglA:$(MglB)_2$ superimposed on the solved structure of MglAGTPγS:$(MglB)_2$ (pdb ID code: 6izw [3]). The AlphaFold model is in light green and the solved structures of MglAGTPγS and of $(MglB)_2$ in cyan and red, respectively. The green sphere indicates $Mg^{2+}$. AlphaFold model rank 1 is shown.

**c.** AlphaFold-Multimer model of MglA:RomY. Left panel, MglA is in yellow, the N-terminal domain of RomY in light teal, and the remainder of RomY in cyan. Middle panel, as in left panel except that only the N-terminal domain of RomY is shown. Right panel, superimposition of MglA from the model of the MglA:RomY complex with the solved structure of MglAGTPγS in light green (pdb ID code: 6h17 [4]). In the modeled structure of MglA, the P-loop is in purple, switch-1 in blue and switch-2 in green. AlphaFold model rank 3 is shown.

**d.** AlphaFold-Multimer model of $(MglB)_2$:RomY. Right panel, the MglB homodimer is in red and the N-terminal domain of RomY in light teal, and the remainder of RomY in cyan. Middle panel, as in left panel except that only the N-terminal domain of RomY is shown. Right panel, superimposition of the MglB homodimer from the model of the $(MglB)_2$:RomY complex with the solved structure of the MglB homodimer in light green (pdb ID code: 6hjm [4]). AlphaFold model rank 1 is shown.

**e.** Alphafold-Multimer model of MglA:$(MglB)_2$:RomY complex. Left panel, the proteins are colored as in **c-d**. Right panel, superimposition of MglA:$(MglB)_2$ from the MglA:$(MglB)_2$:RomY complex with the solved structure of MglAGTPγS:$(MglB)_2$ (pdb ID code: 6izw [3]). The solved structure is in light green and the AlphaFold model in yellow and red. AlphaFold model rank 3 is shown.

pLLDT and pAE for selected rank model of RomY, MglA:$(MglB)_2$, MglA:RomY, $(MglB)_2$:RomY and MglA:$(MglB)_2$:RomY and coordinates of the AlphaFold models are provided in the Source Data file.

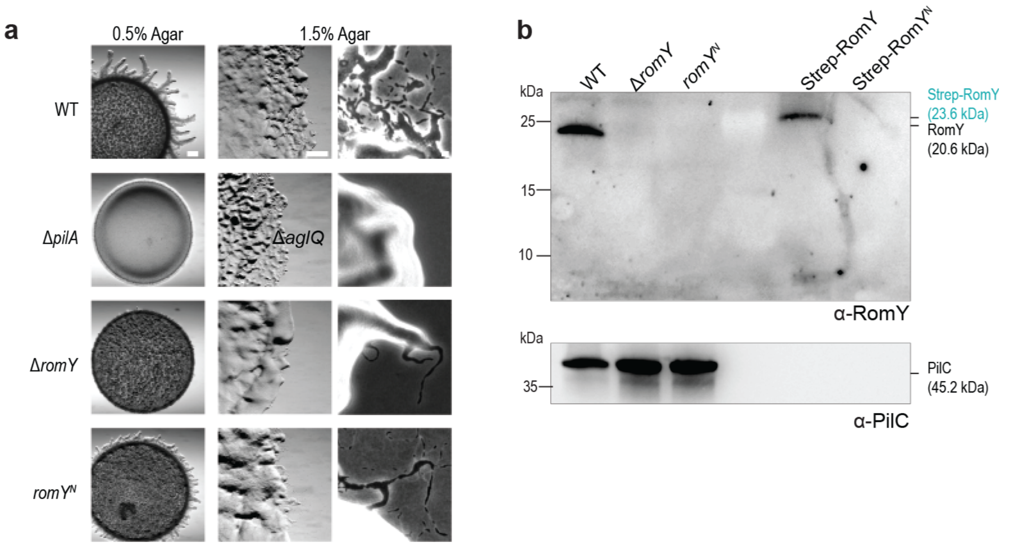


**Figure S5.** The N-terminal domain of RomY has partial RomY activity.

**a.** The N-terminal domain of RomY has partial RomY activity. Cells were incubated on 0.5/1.5% agar with 0.5% CTT to score T4P-dependent/gliding motility. Scale bars, 1mm (left), 500 µm (middle), 50 µm (right). Data are shown from a representative experiment. The experiment was repeated twice with similar results.

**b.** Accumulation of RomY variants. Immunoblot analysis of RomY accumulation. Cell lysates were prepared from same number of cells, separated by SDS–PAGE and probed with α-RomY antibodies and α-PilC antibodies after stripping (loading control). In the two rightmost lanes, 6.0ng and 3.3ng, respectively of purified Strep-RomY and Strep-RomY$^N$ was loaded. The experiment was repeated twice with similar results. RomY$^N$ and Strep-RomY$^N$ have calculated molecular masses of 10.1 and 13.1kDa, respectively.

Source data for **b** are provided in Source Data file.

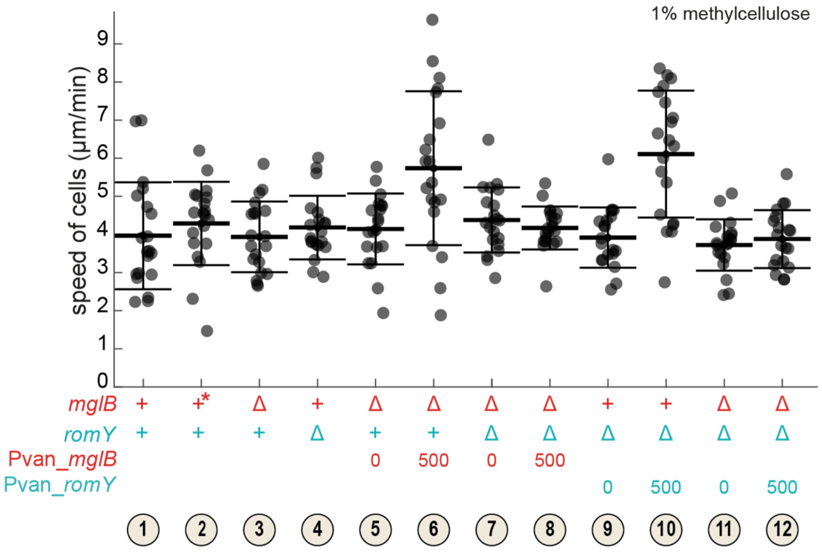


**Figure S6. Overproduction of MglB or RomY does not affect speed of cells moving by T4P-dependent motility.**

Cells were treated as in Figure 4**a** and then T4P-dependent single cell motility analyzed. Strains are numbered as in Figure 4**a**. In the legend, + indicates presence of WT gene, Δ in-frame deletion, 0/500 µM vanillate concentration, and * the WT grown in the presence of 500µM vanillate. Individual data points from a representative experiment with *n* =20 cells are plotted in gray. Because the experiment relies on induction of gene expression, protein levels vary slightly between experiments, making the direct comparison between biological replicates difficult. Consequently, data from only one representative experiment is shown. The cells analyzed are the same as in Figure 4**b**. Boxplots are as in Figure 1**d**. The experiment was repeated twice with overall similar results.

Source data for are provided in Source Data file.

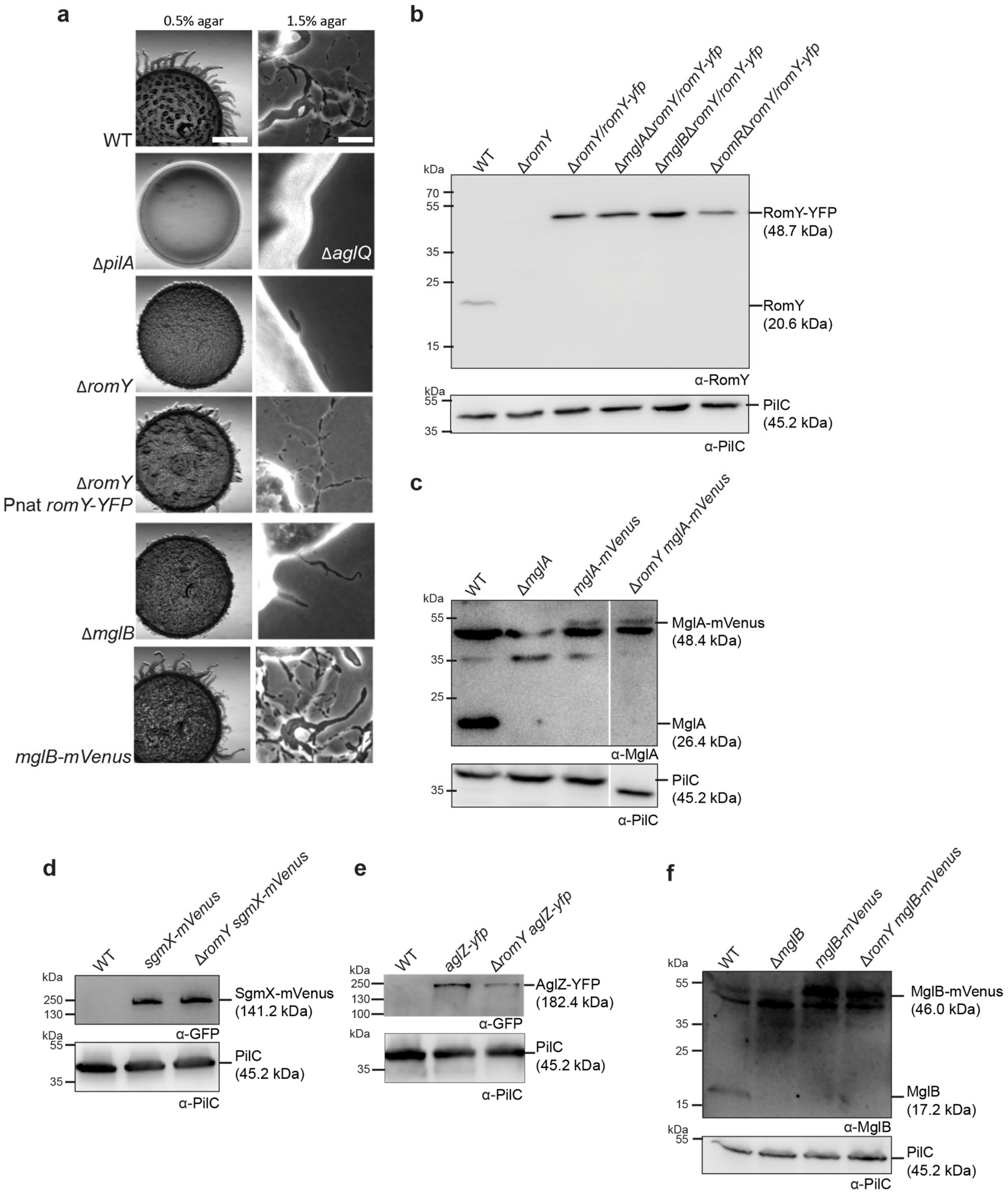


**Figure 7. Analysis of RomY-YFP and MglB-mVenus fusions and accumulate at WT level.**
**a**. RomY-YFP and MglB-mVenus are functional fusions. Motility assays were done as in Figure 1**b**. Scale bars, 1mm (0.5% agar) and 50µm (1.5% agar).
**b**. RomY-YFP accumulation. Immunoblot analysis was done as in Figure 1**c**. RomY and RomY-YFP with calculated molecular masses are indicated. PilC served as a loading control.

**c, d, e, f**. MglA-mVenus, SgmX-mVenus AglZ-YFP and MglB-mVenus accumulation. Immunoblot analysis was done as in Figure 1**c**. Relevant proteins with their with calculated molecular masses are indicated. PilC served as a loading control.
Source data for **b-f** are provided in Source Data file.

**Supplementary Table 1.** Genomic distribution of *mglA*, *mglB*, *romR*, *romX* and *romY* in a set of 1611 prokaryotic genomes

| Species name | MglA | MglB | RomR | RomX | RomY |
|---|---|---|---|---|---|
| Thermovibrio ammonificans HB-1 | Theam_1390 | Theam_1391 | Theam_0615 | Theam_0309 | THEAM_RS03235 |
| Desulfurobacterium thermolithotrophum | Dester_1068 | Dester_1069 | Dester_0564 | Dester_0989 | Dester_0644 |
| Bdellovibrio bacteriovorus HD100 | Bd3734 | | Bd2761 | | Bd0164 |
| Bacteriovorax marinus SJ | BMS_0054 | | BMS_3223 | | |
| Chloracidobacterium thermophilum | Cabther_A0781 | Cabther_A0780 | Cabther_A0541 | Cabther_A1043<br>Cabther_A1145 | Cabther_A0581 |
| Candidatus Koribacter versatilis | Acid345_0708 | | Acid345_0072 | Acid345_2692 | Acid345_1316 |
| Candidatus Solibacter usitatus | | | Acid_3193 | Acid_2709 | |
| Flexistipes sinusarabici | Flexsi_0290 | Flexsi_0291 | Flexsi_0107 | Flexsi_1005 | Flexsi_1362 |
| Calditerrivibrio nitroreducens | Calni_1376 | Calni_1375 | Calni_0658 | Calni_1309 | Calni_0969 |
| Deferribacter desulfuricans | DEFDS_0024 | DEFDS_0025 | DEFDS_0166 | DEFDS_0505 | DEFDS_RS06845 |
| Denitrovibrio acetiphilus DSM 12809 | | | Dacet_3018 | | |
| Geobacter bemidjiensis | Gbem_3962 | Gbem_3963 | Gbem_1530 | Gbem_0745<br>Gbem_3395 | Gbem_0278 |
| Geobacter sp. M21 | GM21_4048 | GM21_4049 | GM21_2686 | GM21_0760<br>GM21_3458 | GM21_0263 |
| Geobacter sp. M18 | GM18_4429 | GM18_4430 | GM18_1365 | GM18_3605<br>GM18_0704 | GM18_0315 |
| Geobacter daltonii FRC-32 | Geob_0373 | Geob_0374 | Geob_3083 | Geob_2164<br>Geob_1026 | Geob_1366 |
| Geobacter uraniireducens | Gura_4332 | Gura_4331 | Gura_1798 | Gura_3141<br>Gura_0880 | Gura_0490 |
| Geobacter metallireducens | Gmet_3417 | Gmet_3418 | Gmet_0955 | Gmet_3436 | Gmet_0550 |
| Geobacter sulfurreducens PCA | GSU_0099 | GSU_0098 | GSU_2046 | GSU_2217<br>GSU_0081 | GSU_3619 |
| Geobacter lovleyi | Glov_3124 | Glov_3123 | Glov_2486 | Glov_0483 | Glov_2834 |
| Pelobacter propionicus DSM_2379 | Ppro_2940 | Ppro_2941 | Ppro_0713 | Ppro_3197 | Ppro_1414 |
| Pelobacter carbinolicus DSM_2380 | Pcar_0379 | Pcar_0378 | Pcar_2316 | Pcar_2949 | Pcar_2721 |
| Sorangium cellulosum So ce56 | sce7249 | sce7248 | sce0012 | sce5932 | sce6926 |
| Haliangium ochraceum | Hoch_6866 | Hoch_6867 | Hoch_3476 | Hoch_2466 | Hoch_6810 |
| Corallococcus coralloides | COCOR_01949 | COCOR_01950 | COCOR_03318 | COCOR_03217 | COCOR_06242 |
| Myxococcus fulvus | LILAB_17450 | LILAB_17455 | LILAB_30440 | LILAB_24525 | LILAB_36235 |
| Myxococcus xanthus DK1622 | MXAN_1925 | MXAN_1926 | MXAN_4461 | MXAN_3350 | MXAN_5749 |
| Stigmatella aurantiaca | STAUR_2690 | STAUR_2691 | STAUR_4816 | STAUR_3813 | STAUR_6415 |
| Anaeromyxobacter sp. Fw109-5 | Anae109_3758 | Anae109_3759 | Anae109_1476 | Anae109_1957 | Anae109_0709 |
| Anaeromyxobacter sp. K | AnaeK_3691 | AnaeK_3692 | AnaeK_1473 | AnaeK_1977 | AnaeK_0709 |
| Anaeromyxobacter dehalogenans 2CP-1 | A2cp1_3774 | A2cp1_3775 | A2cp1_1568 | A2cp1_2062 | A2cp1_0700 |
| Anaeromyxobacter dehalogenans 2CP-C | Adeh_3633 | Adeh_3634 | Adeh_2391 | Adeh_1902 | Adeh_0665 |
| Gemmatimonas_aurantiaca_T_27 | GAU_1116 | GAU_1115 | | GAU_1577 | |

**Supplementary Table 2.** *M. xanthus* strains used in this work

| Strain | Genotype | Source or reference |
|---|---|---|
| DK1622 | Wild type | [5] |
| DK10410 | Δ*pilA* | [6] |
| SA5293 | Δ*aglQ* | [7] |
| SA5958 | Δ*romY* | This work |
| SA6920 | Δ*romY* $P_{nat}$_*romY* (*attB::*pDSZ36) | This work |
| SA4420 | Δ*mglA* | [8] |
| SA3387 | Δ*mglB* | [9] |
| SA3300 | Δ*romR* | [10] |
| SA3683 | Δ*romX* | [11] |
| SA3626 | Δ*mglA* Δ*romY* | This work |
| SA3630 | Δ*mglB* Δ*romY* | This work |
| SA3621 | Δ*romR* Δ*romY* | This work |
| SA5792 | Δ*romX* Δ*romY* | This work |
| SA3936 | Δ*mglB* Δ*romR* | [10] |
| SA3615 | Δ*mglB* Δ*romX* | [11] |
| SA8802 | Δ*frzE* | [11] |
| SA8193 | Δ*frzE* Δ*mglB* | This work |
| SA8316 | Δ*frzE* Δ*romY* | This work |
| SA9138 | $romY^{N}$ | This work |
| SA9113 | Δ*mglB* $P_{van}$_*mglB* (*mxan18-19*::pDSZ31) | This work |
| SA9115 | Δ*mglB* Δ*romY* $P_{van}$_*mglB* (*mxan18-19*::pDSZ31) | This work |
| SA9114 | Δ*romY* $P_{van}$_*romY* (*mxan18-19*::pDSZ30) | This work |
| SA9117 | Δ*mglB* Δ*romY* $P_{van}$_*romY* (*mxan18-19*::pDSZ30) | This work |
| SA6901 | Δ*romY* $P_{nat}$_*romY-YFP* (*attB::*pDK132) | This work |
| SA6913 | Δ*mglA* Δ*romY* $P_{nat}$_*romY-YFP* (*attB::*pDK132) | This work |
| SA6903 | Δ*mglB* Δ*romY* $P_{nat}$_*romY-YFP* (*attB::*pDK132) | This work |
| SA6908 | Δ*romR* Δ*romY* $P_{nat}$_*romY-YFP* (*attB::*pDK132) | This work |
| SA8185 | *mglA-mVenus* | [11] |
| SA7577 | Δ*romY mglA-mVenus* | This work |
| SA7195 | *sgmX-mVenus* | [12] |
| SA11049 | Δ*romY SgmX-mVenus* | This work |
| SA3377 | *aglZ::aglZ-yfp* (pSL65) | [9] |
| SA9102 | Δ*romY aglZ::aglZ-yfp* (pSL65) | This work |
| SA10043 | *mglB-mVenus* | This work |
| SA10040 | Δ*romY mglB-mVenus* | This work |

**Supplementary Table 3.** Plasmids used in this work

| Plasmid | Description | Reference |
|---|---|---|
| pSW105 | $P_{pilA}$, *attP*, $Kan^R$ | [13] |
| pSWU30 | $Tet^R$, *attP* | [14] |
| pBJ114 | $Kan^R$, *galK*¸ vector for generating in-frame deletions | [15] |
| pMR3691 | *vanR* $P_{van}$, $Tet^R$ | [16] |
| pASK-IBA15+ | Vector for overexpression of Strep-tagged proteins; $Kan^R$ | IBA Lifesciences GmbH |
| pET45b+ | Vector for overexpression of $His_6$-tagged proteins; $Ap^R$ | Merck Millipore |
| pDK95 | pBJ114; for generation of in-frame deletion of *romY* | This work |
| pDSZ36 | pSWU30; $P_{nat}$_*romY* | This work |
| pES2 | pBJ114; for generation of in-frame deletion of *mglB* | [9] |
| pDSZ35 | pBJ114; for generation of *romY$^N$* | This work |
| pDSZ31 | pMR3691, *mglB*, $Tet^R$ | This work |
| pDSZ30 | pMR3691, *romY*, $Tet^R$ | This work |
| pDK132 | pSW105; $P_{nat}$_*romY-yfp*, $Kan^R$ | This work |
| pLC20 | pBJ114; for *mglA* replacement by *mglA-mVenus* at native site | [11] |
| pAP35 | pBJ114; for *sgmX* replacement by *sgmX-mVenus* at native site | [12] |
| pSL65 | pBJ114; in-frame integration of *aglZ-gfp* at native site; $kan^R$ | [9] |
| pLC58 | pBJ114; for *mglB* replacement by *mglB-mVenus* at native site | This work |
| pTM1 | Overexpression MglA-$His_6$ | [17] |
| pTM2 | Overexpression $His_6$-MglB | [17] |
| pDSZ32 | Overexpression Strep-RomY | This work |
| pDSZ34 | Overexpression Strep-RomY$^N$ | This work |
| pMAL-c6T | Overexpression $His_6$-MalE | NEB |

**Supplementary Table 4.** Primers used in this work

| Primer | Sequence (5'-3') | Used to construct plasmid |
|---|---|---|
| romYA | ATCGGAAGCTTCCGGAGACGAAGTCCGCGGC | pDK95 |
| romYB | ATCGGTCTAGAGGTGACGGCTTTTTCGAAGGTTTTC | |
| romYC | ATCGGTCTAGAAGTTACCTGGCGGGTGAGGGCG | |
| romYD | ATCGGGAATTCCCACCGTCCGGTGCGGCAGCA | |
| romY_fw | ATCGGGAATTCGCACCGGCATCTCCGTGGAG | pDSZ36, pDK132 |
| romY_rv_stop | ATCGGAAGCTTTCACTGCTCGCCCTCACCCGCCAGGTAA | pDSZ36 |
| romY_yfp_rv | ATCGGAAGCTTTCACTGCTCGCCCTCACCCGCCAGGTAA | pDK132 |
| Yfp fw | ATCGGGGATCCATGGTGAGCAAGGGCGAGGAGCT | |
| Yfp rv stop | ATCGAAGCTTTTACTTGTACAGCTCGTCCATGCC | |
| romY_n-term_A | ATGCGGATCCCCCACGGTGATGCAGTAGTAC | pDSZ35 |
| romY_n-term_B | CCGCGTGGGCGGCTAGTCGTACACCCCGTTGATG | |
| romY_n-term_C | CATCAACGGGGTGTACGACTAGCCGCCCACGCGG | |
| romY_n-term_D | ATGCAAGCTTATGCCCGCGGAGAACAA | |
| DSZ38 | ATCGCATATGATGGGCACGCAACTGGTGATGTACG | pDSZ31 |
| DSZ39 | ATCGGAATTCTTACTCGCTGAAGAGGTTGTCGATATCGTCGTC | |
| DSZ36 | ATCGCATATGACGAAAACCTTCGAAAAAGCCG | pDSZ30 |
| DSZ37 | ATCGGAATTCCTACTGCTCGCCCTCACCCGCCAGGTAA | |
| MglB FW HindIII | ATCGAAGCTTATGGGCACGCAACTGGTGATG | pLC58 |
| mglb_rv_linker | GGAGCCGCCGCCGCCCTCGCTGAAGAGGTTGTC | |
| mglb_ds_fw | ATCGTCTAGACCCGGGAAGCCATGTCCTTCA | |
| MglA_RV_EcoRI | ATCGGAATTCTCAACCACCCTTCTTGA | |
| VenusFWMgla | GGCGGCGGCGGCTCCATGGTGAGCAAGGGCGAG | |
| VenusRV | ATCGTCTAGATTACTTGTACAGCTCGTCCATGCC | |
| DSZ40 | ATCGGAATTCATGACGAAAACCTTCGAAAAAGCCG | pDSZ32, pDSZ34 |
| DSZ41 | ATCGGGATCCCTACTGCTCGCCCTCACCCGCCAGGTAA | pDSZ32 |
| DSZ42 | ATCGGGATCCCTAGTCGTACACCCCGTTGATGAGGTTG | pDSZ34 |

**Supplementary References**


1 Letunic, I. & Bork, P. 20 years of the SMART protein domain annotation resource. *Nucl. Acids Res.* **46**, D493-D496 (2017).

2 Li, W. *et al.* The EMBL-EBI bioinformatics web and programmatic tools framework. *Nucleic Acids Res* **43**, W580-584 (2015).

3 Baranwal, J. *et al.* Allosteric regulation of a prokaryotic small Ras-like GTPase contributes to cell polarity oscillations in bacterial motility. *PLOS Biol.* **17**, e3000459 (2019).

4 Galicia, C. *et al.* MglA functions as a three-state GTPase to control movement reversals of *Myxococcus xanthus*. *Nat. Comm.* **10**, 5300 (2019).

5 Kaiser, D. Social gliding is correlated with the presence of pili in *Myxococcus xanthus*. *Proc. Natl. Acad. Sci. USA* **76**, 5952-5956 (1979).

6 Wu, S. S. & Kaiser, D. Markerless deletions of *pil* genes in *Myxococcus xanthus* generated by counterselection with the *Bacillus subtilis sacB* gene. *J. Bacteriol.* **178**, 5817-5821 (1996).

7 Jakobczak, B., Keilberg, D., Wuichet, K. & Søgaard-Andersen, L. Contact- and protein transfer-dependent stimulation of assembly of the gliding motility machinery in *Myxococcus xanthus*. *PLOS Genet* **11**, e1005341 (2015).

8 Miertzschke, M. *et al.* Structural analysis of the Ras-like G protein MglA and its cognate GAP MglB and implications for bacterial polarity. *EMBO J.* **30**, 4185-4197 (2011).

9 Leonardy, S. *et al.* Regulation of dynamic polarity switching in bacteria by a Ras-like G-protein and its cognate GAP. *EMBO J.* **29**, 2276-2289 (2010).

10 Keilberg, D., Wuichet, K., Drescher, F. & Søgaard-Andersen, L. A response regulator interfaces between the Frz chemosensory system and the MglA/MglB GTPase/GAP module to regulate polarity in *Myxococcus xanthus*. *PLOS Genet.* **8**, e1002951 (2012).

11 Szadkowski, D. *et al.* Spatial control of the GTPase MglA by localized RomR/RomX GEF and MglB GAP activities enables *Myxococcus xanthus* motility. *Nat. Microbiol.* **4**, 1344-1355 (2019).

12 Potapova, A., Carreira, L. A. M. & Søgaard-Andersen, L. The small GTPase MglA together with the TPR domain protein SgmX stimulates type IV pili formation in *M. xanthus*. *Proc. Natl. Acad Sci. USA* **117**, 23859-23868 (2020).

13 Jakovljevic, V., Leonardy, S., Hoppert, M. & Søgaard-Andersen, L. PilB and PilT are ATPases acting antagonistically in type IV pili function in *Myxococcus xanthus*. *J. Bacteriol.* **190**, 2411-2421 (2008).

14 Wu, S. S. & Kaiser, D. Regulation of expression of the *pilA* gene in *Myxococcus xanthus*. *J. Bacteriol.* **179**, 7748-7758 (1997).

15 Julien, B., Kaiser, A. D. & Garza, A. Spatial control of cell differentiation in *Myxococcus xanthus*. *Proc. Natl. Acad. Sci. USA* **97**, 9098-9103 (2000).

16 Iniesta, A. A., García-Heras, F., Abellón-Ruiz, J., Gallego-García, A. & Elías-Arnanz, M. Two systems for conditional gene expression in *Myxococcus xanthus* inducible by isopropyl-thiogalactopyranoside or vanillate. *J. Bacteriol.* **194**, 5875-5885 (2012).

17 Zhang, Y., Franco, M., Ducret, A. & Mignot, T. A bacterial Ras-like small GTP-binding protein and its cognate GAP establish a dynamic spatial polarity axis to control directed motility. *PLOS Biol* **8**, e1000430 (2010).
